# Engineered ACE2-Fc counters murine lethal SARS-CoV-2 infection through direct neutralization and Fc-effector activities

**DOI:** 10.1101/2021.11.24.469776

**Authors:** Yaozong Chen, Lulu Sun, Irfan Ullah, Guillaume Beaudoin-Bussières, Sai Priya Anand, Andrew P. Hederman, William D. Tolbert, Rebekah Sherburn, Dung N. Nguyen, Lorie Marchitto, Shilei Ding, Di Wu, Yuhong Luo, Suneetha Gottumukkala, Sean Moran, Priti Kumar, Grzegorz Piszczek, Walther Mothes, Margaret E. Ackerman, Andrés Finzi, Pradeep D. Uchil, Frank J. Gonzalez, Marzena Pazgier

## Abstract

Soluble Angiotensin-Converting Enzyme 2 (ACE2) constitutes an attractive antiviral capable of targeting a wide range of coronaviruses utilizing ACE2 as their receptor. Here, using structure-guided approaches, we developed divalent ACE2 molecules by grafting the extracellular ACE2-domain onto a human IgG1 or IgG3 (ACE2-Fc). These ACE2-Fcs harbor structurally validated mutations that enhance spike (S) binding and remove angiotensin enzymatic activity. The lead variant bound tightly to S, mediated *in vitro* neutralization of SARS-CoV-2 variants of concern (VOCs) with sub-nanomolar IC_50_ and was capable of robust Fc-effector functions, including antibody-dependent-cellular cytotoxicity, phagocytosis and complement deposition. When tested in a stringent K18-hACE2 mouse model, it delayed death or effectively resolved lethal SARS-CoV-2 infection in a prophylactic or therapeutic setting utilizing the combined effect of neutralization and Fc-effector functions. These data confirm the utility of ACE2-Fcs as valuable agents in preventing and eliminating SARS-CoV-2 infection and demonstrate that ACE2-Fc therapeutic activity require Fc-effector functions.

## INTRODUCTION

Severe acute respiratory syndrome coronavirus 2 (SARS-CoV-2), a betacoronavirus closely related to SARS-CoV-1, is the ninth documented coronavirus capable of infecting humans (*1, 2*) and has led to a devastating on-going pandemic, resulting in nearly 5 million deaths (*3*) worldwide since it first emerged in the Chinese city of Wuhan in late 2019. This highly transmissible airborne pathogen is an enveloped virus with a large, single-stranded, positive-sense RNA genome. Since the genetic sequence became available in January 2020, the development of both traditional vaccines (e.g. inactivated virus, recombinant proteins, viral vectors etc.) and novel RNA/DNA strategies has moved at an unprecedented pace (*4*). The world-wide emergency rollout of vaccines clearly aided in the suppression of viral circulation and reduced the risk of severe illnesses; however, continuous viral evolution and the resulting variants of concern (VOCs) have the potential to circumvent immunity conferred by both natural infection and vaccination. In preparation for the inevitable SARS-CoV-2 VOCs and any future potential pandemic or zoonotic spillovers, it is important that additional interventions and therapies effective against the vast natural CoV reservoirs are developed and stockpiled.

A major antigenic site on the SARS-CoV-2 virion surface is the spike trimer (S) which mediates viral-host membrane fusion and subsequent entry via the primary host cell receptor angiotensin-converting enzyme 2 (ACE2) (*5–8*). Viral entry is initiated by specific interaction of the S1 subunit receptor binding domain (RBD) to ACE2, followed by S2-directed membrane fusion (*9–11*). Most neutralizing antibodies (nAbs) elicited through natural infection and vaccination act by disrupting this interaction; however, selection pressure results in viral escape mutations, in many cases generating VOCs with an enhanced ability to bind host receptors (*12–16*).

Full-length ACE2 consists of an N-terminal protease domain (PD, residue 18-615) which directly engages SARS-CoV-2 RBD, a collectrin-like domain (CLD, residue 616-740), a single transmembrane helix (residue 741-765) and a ∼40 amino-acid intracellular C-terminal domain (*17*). ACE2 is an essential zinc-dependent carboxypeptidase and critical regulator of the renin-angiotensin system (RAS). ACE2 PD converts Angiotensin (Ang) II to Ang 1-7, relieving the vasoconstriction, inflammation and oxidative stress effect of Ang II (*18, 19*).

Membrane bound ACE2 is naturally shed from cell membranes and the circulating ACE2 was reported to play a protective role from SARS-CoV-2 infection in women and children (*20*). Recombinant soluble ACE2 decoys were therefore proposed and tested as potential SARS-CoV-2 therapies since the early onset of the COVID19 pandemic (*21–23*). A pilot clinical trial of human recombinant soluble ACE2 (hrsACE2) administered intravenously (0.4 mg/kg) in a severely SARS-CoV-2 infected patient showed rapid viral clearance in sera, followed by nasal cavity and lung clearance at a later time (*24*). Concomitant with the viral load reduction was a profound decrease of Ang II and a proportional increase of the ACE2 products Ang 1-7 and 1-9 in the plasma. Although ACE2 activity is thought to protect from cardiovascular disorders, an ACE2 inactivated mutant, which has demonstrated equivalent binding to SARS-CoV-2 RBD (*25, 26*), offers a potentially safer therapeutic option applicable to wider cohorts without disturbing the RAS balance.

Since monomeric ACE2 binds to SARS-CoV-2 RBD with only moderate affinity (K_D_ ∼20-30 nM), engineered ACE2 derivatives with improved affinity to SARS-CoV-2 were developed as antiviral therapeutics by several approaches, including deep-mutagenesis coupled with flow-cytometry-based screening (*27–29*), computation-aided design and yeast display (*25, 30*), multimerization of ACE2 (*23, 26, 31-34*), de novo design of ACE2-derived miniprotein and peptides (*35*) and ACE2 decorated vesicles (*36*). Recently, the bivalent ACE2-Fc (i.e. ACE2 extracellular domain grafted onto an IgG1 backbone) molecules have gained considerable attention as they are able to bind SARS-CoV-2 S with increased affinity (mostly through increased avidity) and potently neutralize VOCs, including those resistant to common nAbs (*28, 29*). As most currently investigated ACE2-Fc based therapeutic approaches focus on neutralizing activities, the potential of ACE2-Fcs as agents capable of Fc-mediated effector functions, including antibody dependent cellular cytotoxicity (ADCC), cellular phagocytosis (ADCP) and complement deposition (ADCD) is largely unknown and has not been tested *in vitro* or *in vivo* in models of SARS-CoV-2 infection.

Here we employed a structure-guided approach to develop a series of ACE2-Fc variants, using a human IgG1 or IgG3 backbone. Our variants were engineered to have 1) significantly increased affinity to SARS-CoV-2 RBD derived from the Wuhan strain, B.1.1.7 and B.1.351, 2) enhanced affinity for Fcγ receptors involved in Fc-effector mechanisms and 3) mutations to abrogate the angiotensin enzymatic activity of ACE2. All introduced mutations were validated for mechanism at the molecular level by structural biology approaches. Our best variant cross-neutralized seven SARS-CoV-2 VOCs in a pseudovirus assay with an inhibitory potency comparable to the broad and potent anti-SARS-CoV-2 nAbs and was able mediate an array of Fc-effector activities (**Fig. 1**). When tested in humanized K18-hACE2 mice under prophylaxis or in a single-dose therapeutic setting, the lead ACE2-Fc variant prevented or delayed the lethal SARS-CoV-2 infection. In both cases protection was dependent on the combined effects of direct neutralization and Fc-effector functions.

**Fig. 1.**
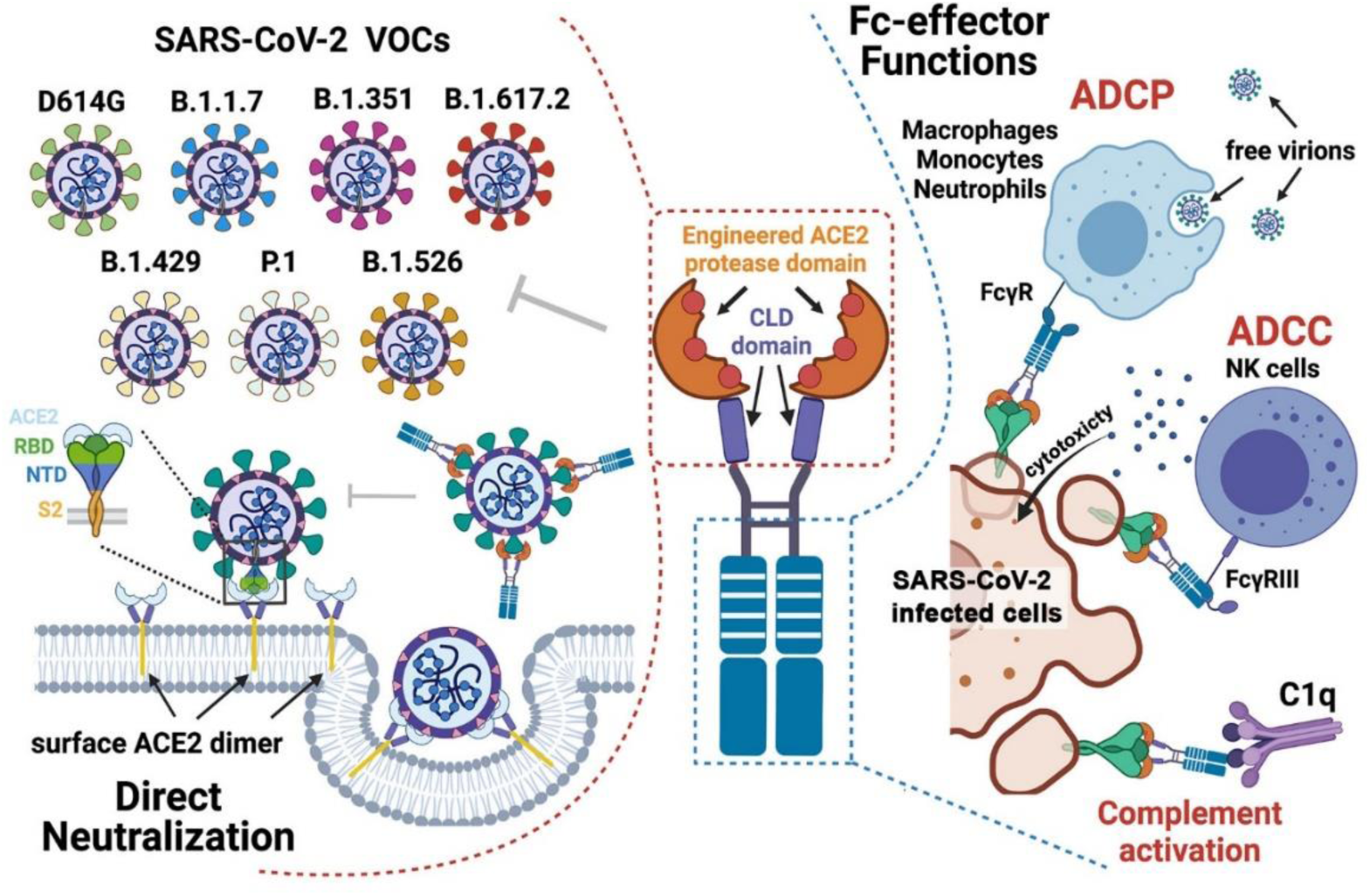
Combined mechanism of direct neutralization and Fc-effector functions by engineered ACE2-Fcs. The figure was generated in *BioRender*.

## RESULTS

### Structure-based engineering of ACE2-Fc variants with enhanced binding affinities to SARS-CoV-2 RBD

A series of hybrid molecules (ACE2-Fc fusion proteins) were generated by replacing the antigen binding fragment (Fab) of human IgG1 or IgG3 with the ACE2 PD (residues 18-615 of ACE2, hereafter referred to as the ACE2_615_ variant) or both the ACE2 PD and CLD (the ACE2 dimerization domain) (residues 18-740 of ACE2, referred to as the ACE2_740_ variant) (**Fig. 2A**). In addition, these ACE2-Fc variants were engineered to contain mutations to 1) increase affinity for the SARS-CoV-2 RBD, 2) abrogate the angiotensin cleavage activity of ACE2 and 3) enhance the affinity for Fcγ receptors involved in Fc-mediated effector mechanisms. Structure-based design was used to identify the ACE2 mutation sites with the potential to increase affinity for the spike RBD binding motif (**Fig. 2B**). We started by analyzing the receptor-antigen interface of two high-resolution ACE2-RBD structures (6M0J (*37*) and 6VW1 (*38*)) systematically to identify interface contacts that could be strengthened and/or optimized. Key interactions important for interface stability, e.g., hydrogen-bonds with distance < 3.0 Å or salt-bridges, were excluded from the design process. Interface residues were then analyzed based on their electrostatic potentials, and ACE2 point mutations that had the potential to enhance charge-complementarity with the RBD were introduced (e.g., K31R, L45D, **Fig. 2B**). Hydrophobic contacts within the flexible regions were also re-designed to improve binary packing and reduce steric repulsion (e.g., F28S), fill empty cavities (e.g., L79F) or improve aromatic interactions (e.g., M82Y). Point mutations to possibly facilitate the hydrogen-bonding network (Q325Y) were also introduced. A list of the ACE2-Fc variants that were generated, expressed, and purified to homogeneity is shown in **Fig. 2C**; the size exclusion chromatographic (SEC) profiles are shown in **Fig. S1**. The enzymatically inactive ACE2-Fc variant was generated by introducing mutations to two Zn^2+^ binding histidines (H374A and H378A, **Fig. 2B**). In addition, to enhance binding to Fcγ receptors present on the effector cell surface and increase Fc-mediated effector functions including ADCC, ADCP and ADCD, the GASDALIE (G236A/S239D/A330L/I332E) mutations (*39–41*) were added to the Fc region of the best performing variants generated with the human IgG1 backbone (**Fig. 2A** and **C**). The best performing ACE2 variant was also fused to the human IgG3 Fc to test if an equivalent IgG3 isotype would display greater Fc-effector activity, as observed for some HIV nAbs (*42, 43*). Finally, an ‘Fc-effector-null’ (L234A/L235A, LALA) mutant (*44*) was generated from the best performing variant to assess the contribution of Fc-effector functions to antiviral activity.

**Fig. 2.**
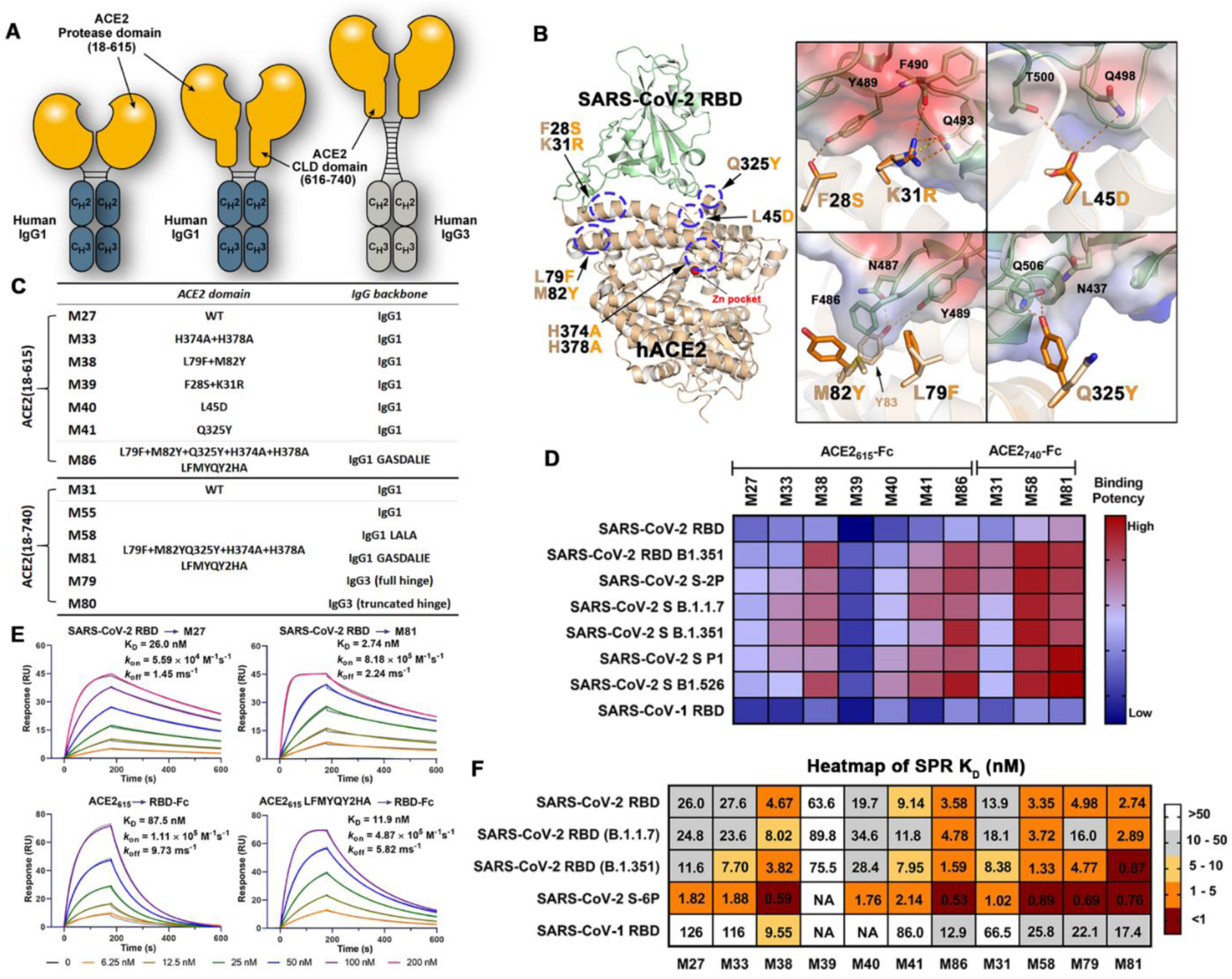
Structure-based development of ACE2-Fc variants with enhanced SARS-CoV-2 RBD affinity. (**A**) Schematic overview of bivalent engineered ACE2-IgG-Fc chimeras (**B**) The structure-based approach for the prediction of ACE2 mutations with the potential to improve the SARS-CoV-2 RBD binding affinity. The 2.45 Å crystal structure of the ACE2 (light yellow) and RBD (pale green) complex (RDB: 6M0J) used for interface residue analysis (left panel) with residues selected for mutation shown as sticks. Blow-up views of the mutation sites with wild type and mutated residues shown as sticks (right panel). The RBD is shown as a semi-transparent electrostatic potential surface (red for negatively charged residues and blue for positively charged residues) with a green cartoon for the polypeptide backbone. Wild-type and mutated ACE2 residues are colored as pale-yellow and orange respectively. (**C**) A list of the developed ACE2-Fc variants. (**D**) A heat map showing the binding efficacy of ACE2-Fc variants to SARS-CoV-2 RBD_wt_, RBD_B.1.351_, SARS-CoV RBD and the selected SARS-CoV-2 VOCs. Binding was measured by ELISA using SARS-CoV-2 antigens immobilized on the plate and ACE2-Fc in the concentration range of 0.05-125 nM. Area-under-curve (AUC) for the unsaturated binding region (0.05-2.5 nM, Fig. S2) were calculated and plotted as heatmap. (**E**) SPR-based kinetic measurement of SARS-CoV-2 RBD_wt_ binding to immobilized M27 or M81 (top panel), and monomeric wtACE2_615_ or ACE2_615_(LFMYQY2HA) to immobilized SARS-CoV-2 RBD_wt_-Fc (bottom panel). Experimental data are shown as colored curves overlapped with the 1:1 Langmuir fitting model in grey. (**F**) The dissociation constants (K_D_) for SARS-CoV-2 and SARS-CoV antigens binding to ACE2-Fc variants as measured by SPR. ACE2-Fc variants were immobilized on Protein A chip and various viral antigens including SARS-CoV-2 RBD_wt_, RBD_B.1.1.7_, RBD_B.1.351_, S-6P and SARS-CoV-1 RBD were injected as flow analytes. The K_D_ values were determined using 1:1 Langmuir model. The experimental binding curves and the detailed kinetic constants are shown in **Fig. S3** and summarized in **Table S1**).

The initial screening of ACE2-Fc variant binding affinity to SARS-CoV-2 wild-type (wt, Wuhan-Hu-1 strain) RBD and selected VOC RBDs (e.g., B.1.1.7 and B.1.351) were performed by ELISA (**Fig. 2D** and **S2**) and surface plasmon resonance (SPR) (**Fig. 2F** and **S3, Table S1**). The wt ACE2_615_-Fc (M27, **Fig. 2C**) bound to RBD_wt_ with a dissociation constant (K_D_) of 26 nM, consistent with reported data (*27, 45*), and around 5-times higher than the affinity of SARS-CoV-1 RBD binding. A slight enhancement (1.4-2.0-fold) to the binding affinity of all RBDs tested was observed for the ACE2_740_-Fc (M31) variant over the shorter ACE2_615_-Fc (M27) variant (**Fig. 2F, S2-3**). Furthermore, the ACE2_615_-Fc variant with H374A/H378A mutations (M33, **Fig. 2C**) displayed RBD binding comparable to the wild-type (M27), indicating that the zinc-site disrupting substitutions do not interfere with the ACE2-RBD binding interface.

Among the interface mutations that could potentially facilitate RBD binding (**Fig. 2D** and **F**), the dual L79F/M82Y mutant (M38 in **Fig. 2C**) and the single Q325Y mutant (M41) showed a 3-6-fold and 1.4-2.8-fold enhancement in binding affinity to the RBD, respectively, compared to the unmodified ACE2_615_-Fc (M27). In contrast, variants with F28S/K31R mutations (M39) or an L45D mutation (M40) showed reduced or unchanged binding affinity to SARS-CoV-2 RBDs. Interestingly, combining the enhancing mutations (L79F/M82Y and Q325Y) generated variants with significantly increased affinity for RBD_wt_. Variants with the combined L79F/M82Y/Q325Y and H374A/H378A mutations (referred to as LFMYQY2HA mutation) were fused to GASDALIE IgG1 Fc (M81 and M86), LALA IgG1 Fc (M58) or IgG3 Fc (M79 and M80). As shown in **Fig. 2D** and **F**, the best-performing variant, M81, showed increased affinity compared to the unmodified ACE2_615_-Fc (M27) by ∼8.5-13.3-fold to RBD_wt_, RBD_B.1.1.7,_ and RBD_B1.351_ (K_D_ range of 0.87-2.89 nM). This binding enhancement is likely a result of the faster RBD association (*k*_on_) (5-14-fold) as the dissociation constant (*k*_off_) was similar to wild-type levels (**Fig. 2E**). We observed decreased binding affinity for ACE2 variants tested in the SPR format where SARS-CoV-2 RBD-Fc fusion was immobilized, probably due to the slower tumbling rate of monomeric ACE2_615_ (∼75kD) when acting as the soluble analyte compared to the smaller RBD (∼26 kD) (**Fig. 2E**). Taken together, our lead ACE2-Fc variant M81 showed comparable RBD affinity to the reported best-in-class engineered ACE2-Fcs (K_D_ below 1 nM) (*25, 27–29*) and to many neutralizing antibodies isolated from SARS-CoV-2 patients.

Of note, the ACE2_740_-Fc showed significantly enhanced binding affinity as compared to ACE_615_-Fc. ACE_740_ grafted onto an IgG3 backbone (M79) bound to the variants tested with noticeably decreased affinity (**Fig. 2F**). These data point towards the possibility that while the extended CLD likely increases the structural plasticity of ACE2-Fc (*46*) and facilitates avid interaction with spike (*25, 47*), the elongated IgG3 hinge most likely restricts the ACE2 mobility required for optimal RBD recognition.

### Molecular basis for the enhanced SARS-CoV-2 S binding and enzymatic inactivation of engineered ACE2-Fc

To dissect the molecular basis of the enhanced affinity for diverse RBDs and abrogated enzymatic activity of our best ACE2 variant, monomeric ACE2_615_ with the LFMYQY2HA mutations was co-crystallized with SARS-CoV-2 RBD and the structure was determined to 3.54 Å resolution. Four ACE2-RBD complexes were presented in the asymmetric unit (ASU) of the crystal and the final model refined to an R_work_/R_free_ of 0.24/0.29 (**Fig. 3** and **S4**, **Table S2**). The overall interface and contact residues of ACE2_615_ LFMYQY2HA-RBD largely resemble the ACE2_wt_-RBD (**Fig. 3B** and **3E**) with a slightly larger total buried surface area (BSA) (957.4 Å^2^) as compared to the BSA of ACE2_wt_-RBD complex (average of 869.1 Å^2^ calculated from the available ACE2_wt_-RBD structures) and that of the only other RBD enhancing ACE2 engineered variant available in the PDB (BSA of 908.9 Å^2^, PDB: 7DMU) (**Fig. S4B**). In the ACE2_wt_-RBD complex, over 70% of the RBD contacts are mediated by the ACE2 α1-helix (residues 18-52) which contains many reported RBD-binding-enhancing mutations (*25, 27, 28*) (**Fig. 3B**). To differentiate the engineered ACE2 mutations reported previously from those identified in this study, we divided the RBD contact surface on ACE2 into five sub-sites, designated: Site-I (residues 18-45), Site-II (residues 79-83), Site-III (residues 324-330), Site-IV (residues 353-357) and Site-V (residues 386-393) (**Fig. 3B**). The introduced affinity enhancing mutants L79F/M82Y and Q325Y map to Site-II and Site-III, respectively, flanking the α1-helix/Site-I at the furthest edge of the RBD contact surface (**Fig. 3C**). In the ACE2_wt_-RBD structure (PDB: 6M0J), the RBD ridge (residues 473-490) is weakly associated with Site-II residues and represents the most mobile segment with the highest B-factors of residues among the interface. Contacts within this region, specifically with RBD ridge residue F486 which interacts with ACE2 residues Y83, L79 and M82, are significantly stabilized in complex with the engineered ACE2_615_ LFMYQY2HA mutant. Specifically, L79F and M82Y are in face-to-face or face-to-edge stacking with F486 in two of the ACE2_615_ LFMYQY2HA-RBD complex copies in ASU of the crystal (assembly A and B) while only L79F is in face-to-edge stacking with F486 in assembly C and D of the ASU (**Fig. 3D**). As a result, residues (G485-F486-N487) of the RBD ridge with better hydrophobic packing in copies A and B have the lowest relative B-factor values, followed by those in copies C and D and in two wt ACE2-RBD crystal structures (**Fig. S4C**). This observation supports the stabilizing effect of the ACE2 L79F/M82Y mutations to the RBD ridge although this effect depends somewhat on how well the three Site-II aromatic residues (L79F, M82Y, and Y83) pack against RBD residue F486. In contrast, the other RBD-affinity-enhancing mutant, Q325Y, in all four copies of the ASU uniformly forms a strong hydrogen bond with RBD residue Q506 which is not involved in the receptor binding in the ACE2_wt_-RBD structure (**Fig. 3D, S4A** and **D**). This additional hydrogen bond is responsible for a 1.4-2.8-fold enhancement to the binding affinity of the mutant to SARS-CoV-2 RBD (**Fig. 2E** and **S3**). Of note, the two anchor RBD residues F486 and Q506, which interact with L79F/M82Y and Q325Y, respectively, are invariant among the SARS-CoV-2 VOCs to date (**Fig. 3E**), suggesting that these introduced RBD-enhancing mutations could be equally effective against these SARS-CoV-2 escape variants (**Fig. S5A-C**) and a wide variety of CoVs that utilize ACE2 as receptors (**Fig. S5D**).

**Fig. 3.**
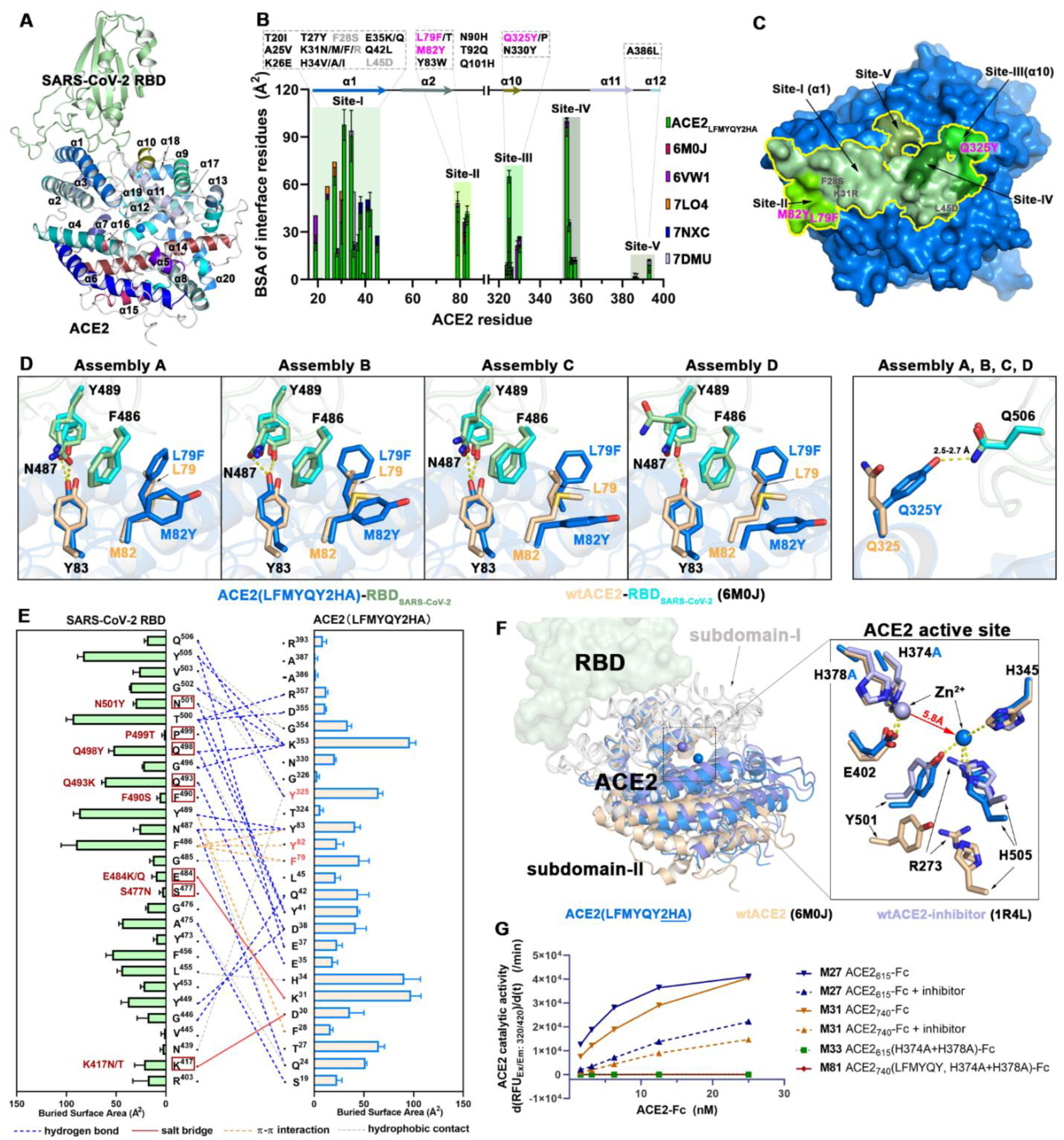
Crystal structure of the protease domain of ACE2_615_ with LFMYQY2HA mutations in complex with the RBD of SARS-CoV-2. (**A**) Overall structure of the ACE2_615_ LFMYQY2HA– RBD complex. ACE2 helices (α1-α20) are colored and labeled according to the criteria defined in (*48*) in which helix α1 serves as the major element of RBD binding. (**B-E**) Properties of the ACE2_615_ LFMYQY2HA–RBD interface. (**B**) Comparison of the buried-surface-area (BSA) of individual ACE2 residues involved in RBD binding between ACE2_615_ LFMYQY2HA, ACE2 wild type (6M0J (*37*), 6VW1 (*38*), 7LO4 (*85*), 7NXC (*86*)), or ACE2 with RBD enhancing mutations as reported by others (7DMU) (*28*). BSA values were calculated by PISA (*72*). The RBD binding residues are classified into Site I (residues 19-45), Site II (residues 79-83), Site III (residues 324-330), Site IV (residues 353-358), and Site V (residues 386-394). ACE2 RBD binding enhancing mutations from this study (pink for enhancing mutations, grey for null mutations) or reported by others are shown (black) above the plot. (**C**) The RBD footprint on ACE2_615_ LFMYQY2HA with details of the overall structure of ACE2. ACE2_615_ LFMYQY2HA is shown as a blue surface with regions that contribute to RBD binding (i.e. BSA>0) contoured by a yellow line. Shades of green are used to color Sites I-IV using the definitions defined in (**B**). (**D**) Molecular details of the interaction of the introduced L79F/M82Y and Q325Y mutations with the RBD. Each individual mutation site was analyzed within the context of the wild-type ACE2 bound to RBD (6M0J) for each of the four individual ACE2_615_ LFMYQY2HA–RBD complexes present in the asymmetric unit of the crystal. L79F/M82Y shows slightly different orientations of introduced side chain within different copies in the asymmetric unit while the side chain of Q325Y is invariant between copies (see also Fig. S4A). Hydrogen bonds (with a distance < 3.5 Å) are depicted as dashed-lines. (**E**) The interaction network at the ACE2_615_ LFMYQY2HA–RBD interface. The antigen-receptor interactions defined by a 5-Å distance criterion cutoff are shown as lines with a diagram of BSA values for individual interface residue shown on the side. Hydrogen bonds and salt bridges (bond lengths < 3.5 Å) are shown as blue dashed lines and red solid lines respectively. Hydrophobic interactions or bond distances between 3.5–5.0 Å are shown as grey dotted lines. π -π interactions (face-to-edge or face-to-face) between aromatic residues are shown as orange broken lines. The ACE2 mutations L79F, M82Y, and Q325Y are highlighted in red and the RBD mutated residues (brown) identified in SARS-CoV-2 VOCs are marked with brown boxes. (**F**) Structural changes introduced by H374A/H378A mutations. ACE2_615_ LFMYQY2HA–RBD, wild type ACE2-RBD, and an inhibitor bound wild type ACE2-RBD (PDB: 1R4L) (PDB:6M0J) are aligned based on the ACE2 subdomain I ( subdomain organization is defined as in (*48*)) A low up view into of the wild type ACE2 active site (right panel). The catalytic zinc ion in the native ACE2 (wheat) is coordinated by H374, H378 and E402 from sub-domain I. In the H374A/H378A mutant, the zinc ion moves to a substrate/inhibitor-binding site ∼5.8 Å from the original zinc binding site and is bound by residues R273, H345, Y501, and H505 (see also **Fig. S4B-C**) which form the substrate binding pocket of the wild type ACE2. (**G**) Angiotensin converting activity of the ACE2-Fc variants. The slopes of the initial linear region of the reaction, as reflected by the fluorometric product formation, were plotted against the indicated ACE2-Fc concentrations.

The second set of mutations, H374A and H378A, were introduced at the zinc-binding site to disrupt angiotensin-converting activity without affecting the ACE2-RBD recognition. Towler *et al*. (*48*) first described the ACE2 PD subdomains I and II that form the active site cleft and revealed the ligand-dependent subdomain-II closure. **Fig. 3F** shows the structural alignment based on the ACE2 domain of the ACE2_615_ LFMYQY2HA-RBD complex, the ACE2_wt_-RBD ‘apo’ complex (6M0J), and ACE2_wt_ with a bound inhibitor (1R4L). There were no significant differences in the overall structure of subdomain-I and the RBD binding regions, consistent with our finding that the H374A and H378A mutations do not interfere with binding to SARS-CoV-2 RBD (**Fig. 2D** and **2F**). However, significant differences were observed in the subdomain-II conformation and the inter-domain Zn^2+^-mediated active site (**Fig. 3F**). ACE2_615_ LFMYQY2HA adopts a subdomain-II conformation that only partially overlaps with either apo or inhibitor-bound, closed conformation of ACE2 with more similarity to the latter. The observed changes can be attributed to the introduced H374A/H378A mutations and relocation of the Zn^2+^ binding site in ACE2_615_ LFMYQY2HA. In ACE2_wt_, the catalytic Zn^2+^ is coordinated by three subdomain I residues H374, H378 and E402, but in the H374A/H378A mutant we found this zinc pocket empty, and a spherical electron density appeared ∼5.8 Å from the original binding site within the substrate/inhibitor binding site (**Fig. S4E-F**). As there were no divalent cations in the crystallization or protein buffers, we attributed this density to endogenous zinc. This new Zn^2+^ ion was coordinated by R273, H345, Y501, and H505 which are responsible for angiotensin substrate recognition in the ACE2_wt_ (**Fig. S4E-F**). We speculate that this alternate zinc coordination site is of structural importance to ACE2 structural integrity but is non-catalytic. A dual-zinc-coordination site is not uncommon in aminopeptidases which remove N-terminal amino acids (*49*). Collectively, our data provides the structural basis for the ACE2 inactivation induced by the zinc-coordination mutations. As predicted, the H374A/H378A mutations were sufficient to abrogate angiotensin converting activity in the ACE2-Fc (**Fig. 3G** and **4G**).

**Fig. 4.**
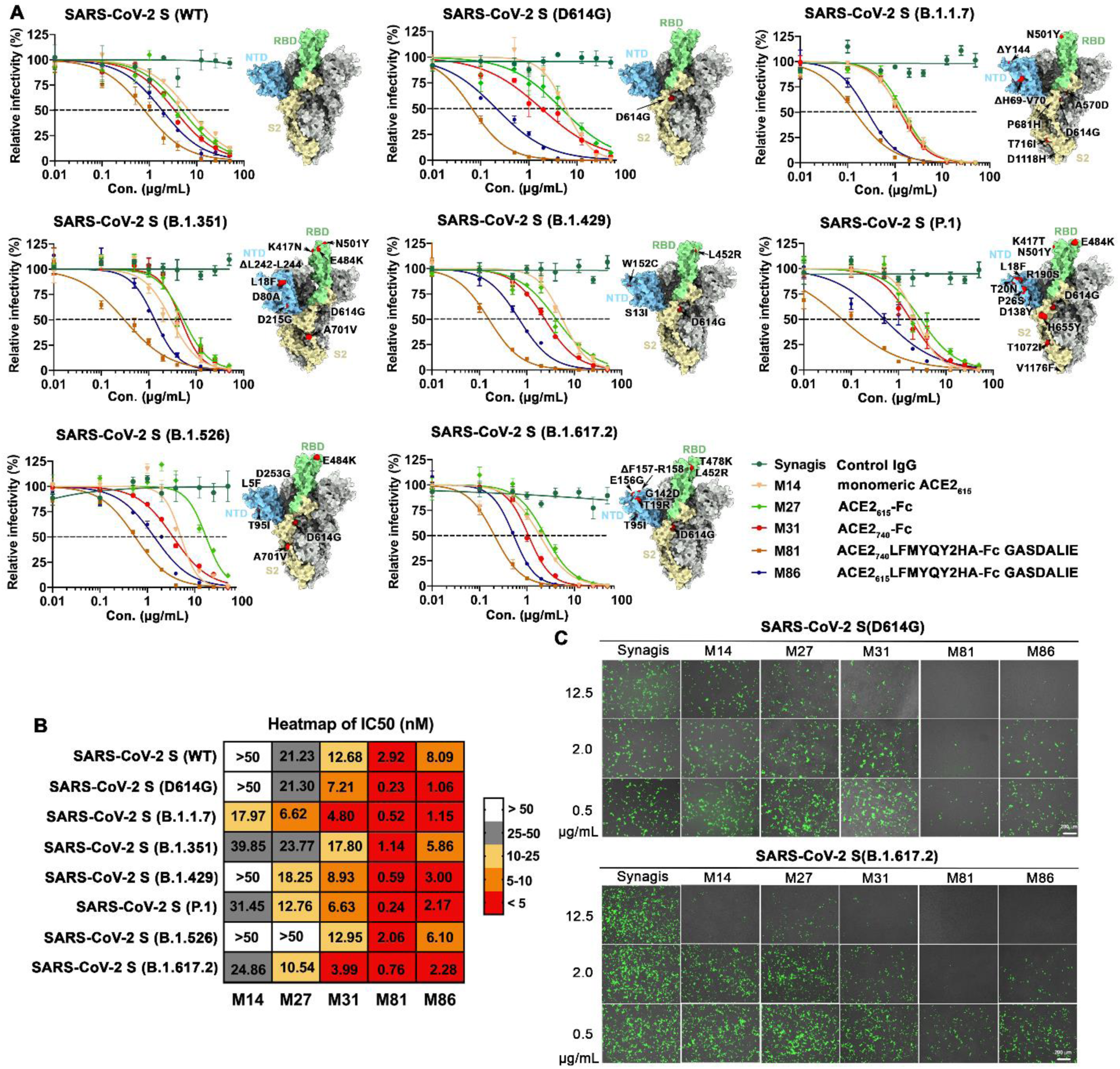
Broad neutralization of engineered ACE2-Fcs against SARS-CoV-2 PsV. (**A**) Dose response neutralization curves of SARS-CoV-2 lentivirus pseudotyped with eight SAR2-CoV-2 S variants. hACE2 expressing 293T cells were infected with different variants of SARS-CoV-2 PsV in the presence of varying concentrations of monomeric ACE2_615_, selected ACE2-Fc variants, Synagis IgG (negative control) or PBS saline. Infectivity was quantified by the cellular luciferase signal 48h post infection. Relative infectivity was normalized by the luciferase signal in infected cells without intervention (PBS saline). The spike graphics for individual VOCs were generated using PDB 7C2L with mutation sites colored in red. Data are shown as mean ± SEM from three independent replicates. (**B**) Heat-map summary of neutralization IC_50_ values for the ACE2-Fc variants tested. (**C**) Representative fluorescent imaging of hACE2-293T cells that were infected with SARS-CoV-2 S (D614G, top) or (B.1.617.2, bottom) in the presence of the indicated concentrations of ACE2-Fc variants. Images are shown as merged bright field (cell shape) and green field (ZsGreen signal). Scale bar: 200 μm. *n* = 3 replicates/group.

### Engineered ACE2-Fcs show potent neutralization of SARS-CoV-2 VOCs *in vitro*

Our engineered ACE2-Fcs showed enhanced affinity for SARS-CoV-2 RBD derived from the Wuhan-Hu-1 strain and several VOCs (**Fig. 2**). To evaluate if the increased affinity for the S glycoprotein translated to improved neutralizing activity, we tested the best performing ACE2 LFMYQY2HA-Fc variants in an *in-vitro* neutralization assay using lentivirus pseudotyped with the spike from eight SARS-CoV-2 strains, including Wuhan-Hu-1 (wt), D614G, B.1.1.7 (Alpha), B.1.351 (Beta), P1 (Gamma), B.1.429 (Epsilon), B1.526 (Iota) and the currently dominant B.1.617 (Delta). Pseudotyped lentiviruses (PsV) carrying the reporter genes of luciferase (Luc2) and ZsGreen-1 were generated as previously described (*50*). hACE2-expressing 293T cells were infected with SARS-CoV-2 PsV (10^6^ RLU) and then pre-incubated for 1h with varying concentrations (0.01-50ug/mL) of wild-type monomeric ACE2_615_ (M14), wild type ACE2-Fc (M27 or M31), or ACE2 LFMYQY2HA-Fc variants (M81 or M86). Quantitative luciferase readout and live cell imaging for ZsGreen were performed 48h post-infection (**Fig. 4** and **S6-7**).

For all the tested SARS-CoV-2 PsVs, neutralization by the bivalent ACE2-Fcs was >2-fold greater than monovalent ACE2_615_ (M14), as reflected in the half-maximal inhibitory concentrations, IC_50_ (molar units, **Fig. 4B**), highlighting the importance of multivalency in soluble ACE2-based therapeutics (*23*). The ACE2_wt_-Fcs (M27 and M31) neutralized SARS-CoV-2 PsV_wt_ with an IC_50_ of 21.2 nM and 12.6 nM, respectively (**Fig. 4A-B**), which concurs with their SPR K_D_ values (26 nM and 13.9 nM, respectively) for SARS-CoV-2 RBD (**Fig. 2F**) and with previous reports (*25–28*).

Both engineered ACE2-Fc variants, M81 (ACE2_740_ LFMYQY2HA-Fc) and its truncated version M86 (ACE2_615_ LFMYQY2HA-Fc), cross-neutralized eight SARS-Cov-2 PsV with low nano-molar IC_50_ values (**Fig. 4A-B**). Of note, the best-performing variant, M81, which bound to SARS-CoV-2 RBD variants with a K_D_ of 0.87-2.89 nM, inhibited SARS-CoV-2 PsV_wt_ with an IC_50_ of 2.92 nM. Interestingly, the CLD-containing M81 showed better neutralization than the CLD-lacking M86, as reflected by a 2.2-9.0-fold reduction of IC_50_ across the tested VOCs. Similar IC_50_ differences in the range of 1.6-3.0-fold were observed between M31 and M27 for VOCs neutralization, which further supports the observation that the collectrin-like domain promotes SARS-CoV-2 S/ACE2-Fc recognition (*25, 46*).

Notably, we observed significantly better neutralization with M81 and M86 toward VOCs variants containing the D614G mutation (**Fig. 4A-C**). As the first recurrent S mutation present in all VOCs to date, D614G has been shown to shift the RBD conformational equilibrium to a wider range of open trimer states, facilitating enhanced receptor binding and virus transmission (*51*). Indeed, the single substitution of D614G substantially enhanced PsV infectivity, as demonstrated by the higher ZsGreen signal (**Fig. 4C** as compared to PsV_wt_ (green fluorescence barely seen, data not shown)). Consistent with a recent study (*52*), this allosteric mutation also makes VOCs more susceptible to RBD-specific nAbs and ACE2-Fc, as shown by the >10-fold reduction in the M81 IC_50_ to PsV_D614G_. Although other SARS-CoV-2 VOCs were less sensitive to M81 neutralization, IC_50_ values were still in the range of 0.24-2.06 nM, comparable to high-affinity antibodies isolated from convalescent patients (*53–55*). Taken together, our PsV-based neutralization studies demonstrated that the best RBD-binder M81 can neutralize SARS-CoV-2 PsV and VOCs that possess D614G with enhanced potency, i.e., with a low nano-molar IC_50_ which is ∼10-90 fold lower than wtACE2_615_-Fc (M27).

### Engineered hACE2-Fc efficiently blocks SARS-CoV-2 PsV transduction in K18-hACE2 mice

Next, we tested the capacity of engineered hACE2-Fc to prevent SARS-CoV-2 viral transduction *in vivo* using an adapted pseudovirus-based mouse infection protocol (*56*) that provides a safe alternative for evaluating antivirals *in vivo* under ABSL-2 conditions (**Fig. 5** and **S8**). Lentivirus pseudotyped with S from two highly infective SARS-CoV-2 variants, D614G and B.1.617, were produced using the same protocol as above (*50*) and concentrated by PEG 8000 in the final step. In K18-hACE2 transgenic mice, 5 μg or 25 μg ACE2-Fc (M27 or M81), or 25 μg Synagis (control IgG) was delivered intranasally (i.n.) 1 h prior to administration of replication-defective SARS-CoV-2 PsV_D614G_ or PsV_B.1.617_ (i.n., ∼1× 10^8^ PFU) expressing Luc2 firefly luciferase. Longitudinal bioluminescence imaging (BLI) on live mice was performed at 4-, 8- and 12-days post infection (dpi) (**Fig. 5A-B**). Due to the non-replicative nature of the pseudovirus and sub-optimal luciferase reporter for in-vivo imaging, BLI signal was only detected around the nasal cavity. In control-IgG treated mice, the fluorescent signal increased by a factor of >1000 over the 12-day time course, clearly demonstrating that SARS-CoV-2 PsV was capable of transducing cells in the nasal cavity of K18-hACE2 mice (**Fig. 5C-E** and **S8E**). The luminescence intensity increased between 0-8 days and reached a plateau thereafter.

**Fig. 5.**
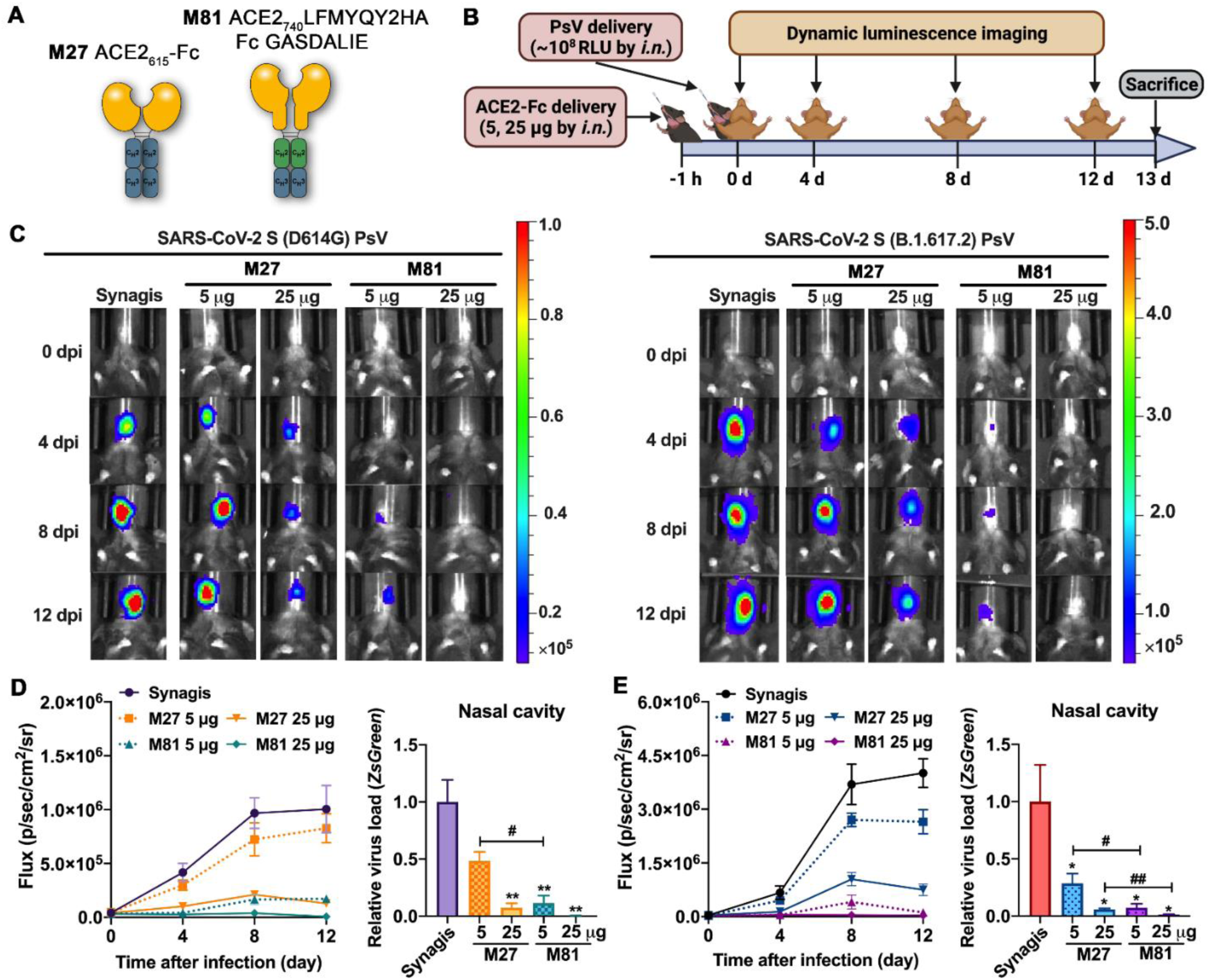
*In vivo* efficacy of engineered ACE2-Fc in blocking SARS-CoV-2 PsV transduction in K18-hACE2 mice. (**A**) Scheme of wt ACE2_615_-Fc (M27) and the engineered variant M81. (**B**) Experimental design of the PsV-challenged K18-hACE2 mouse model. M27, M81 or Synagis (a negative control) were intranasally administrated 1 h before SARS-CoV-2 PsV_D614G_ or PsV_B.1.617.2_ challenge (i.n., ∼10^8^ RFU), and non-invasive luminescence imaging was performed every 4 days. (**C**) Representative BLI images that indicate the luciferase signal for PsV_D614G_ (left) and PsV_B.1.617.2_ (right). (**D-E**) Quantification of luciferase signal as flux (photons/s) computed non-invasively in the nasal area (left) and real-time PCR quantification of SARS-CoV-2 PsV RNA loads (targeting ZsGreen) in the nasal cavity at the end-point (13 dpi, right) for PsV_D614G_ (**E**, n=3-4) and PsV_B.1.617.2_ (**F**, n=4-5). The data are shown as means ± the SEM. Kruskal-Wallis test with Dunn’s post hoc test: \**P*<0.05, \*\**P*<0.01 versus synagis and ^#^*P*<0.05, ^##^*P*<0.01.

Mice pre-treated with 5 μg of wtACE2_615_-Fc (M27) prior to PsV_D614G_ challenge showed a minor reduction in luminescent signal, while increasing M27 to 25 μg led to a viral inhibition of >85% of the control cohort. In contrast, only 5 μg of our lead variant ACE2 LFMYQY2HA-Fc (M81) was sufficient to reach >85% inhibition, while 25 μg M81 nearly completely eradicated the BLI signal (**Fig. 5C-D**). Endpoint analysis (13 dpi) after necropsy to estimate viral transduction levels (*ZsGreen* mRNA level) in the nasal cavity demonstrated that the low-dose M81 had equivalent antiviral activity as high-dose M27.

In the PsV_B.1.617_ challenge model, which demonstrated considerably higher infectivity than PsV_D614G_ (**Fig. 5C**), 5 μg M27 pretreatment failed to inhibit viral transduction and the 25 μg M27 treatment group maintained ∼20% luciferase signal at 12 dpi as compared to the control group (**Fig. 5E**). Conversely, 5 μg of M81effectively protected mice, with only basal luciferase signal detected at 12 dpi. Our SARS-CoV-2 PsV-challenge mouse model thus demonstrated that engineered ACE2-Fc M81 has the potential to effectively inhibit of SARS-CoV-2 infection when administered prophylactically.

### Engineered ACE2-Fc variants mediate potent Fc-effector functions

To dissect the mechanism of action for protection observed in mice we assessed engineered ACE2-Fc variants for Fc-mediated effector functions. Variants were assessed *in vitro* with assays that quantitatively measure the antibody dependent cellular cytotoxicity (ADCC), phagocytosis (ADCP) and complement deposition (ADCD) as previously described (*57–60*). To investigate ADCC potential, T-lymphoid cells with resistance to non-specific NK cell-mediated cell lysis (CEM.NKr) and stable surface expression of SARS-CoV-2 S were mixed at a 1:1 ratio with parental CEM.NKr CCR5+ cells and used as target cells with PBMCs from healthy donors used as effector cells. For the ADCP assay, multiple targets were utilized, including fluorescent microspheres coated with SARS-CoV-2 RBD or S-6P and S-expressing CEM.NKr cells and the human monocytic THP-1 cell line was used as phagocytic effector cells. Similarly, in the ADCD assay, multiplex assay beads coated with SARS-CoV-2 RBD or S-6P were used to form immune complexes with varied concentrations of tested ACE2-Fc variants and the complement C3 deposition was detected by an anti-guinea pig C3 antibody.

As shown in **Fig. 6A**, engineered ACE2 LFMYQY2HA-Fc variants (M58, M81, M86) showed higher surface binding to cell targets expressing SARS-CoV-2 S compared to ACE2-Fcs without RBD-binding enhancing mutations (M27 and M31). We observed also low binding efficiency of M79, ACE2_740_ LFMYQY2HA grafted to the IgG3 backbone, most likely due to the mismatch of the anti-IgG1 secondary Ab used for fluorescent quantification (Materials and methods). Interestingly, although all ACE2-Fcs tested showed variable binding to the target cells (**Fig. 6A**), only variants with Fc GASDALIE (M81 and M86) were able to stimulate robust cytotoxic responses, leading to the killing of ∼40% of target cells expressing S protein (**Fig. 6B**). No ADCC activity was observed for the equivalent ACE2 variants with either Fc LALA or IgG3 backbones.

**Fig. 6.**
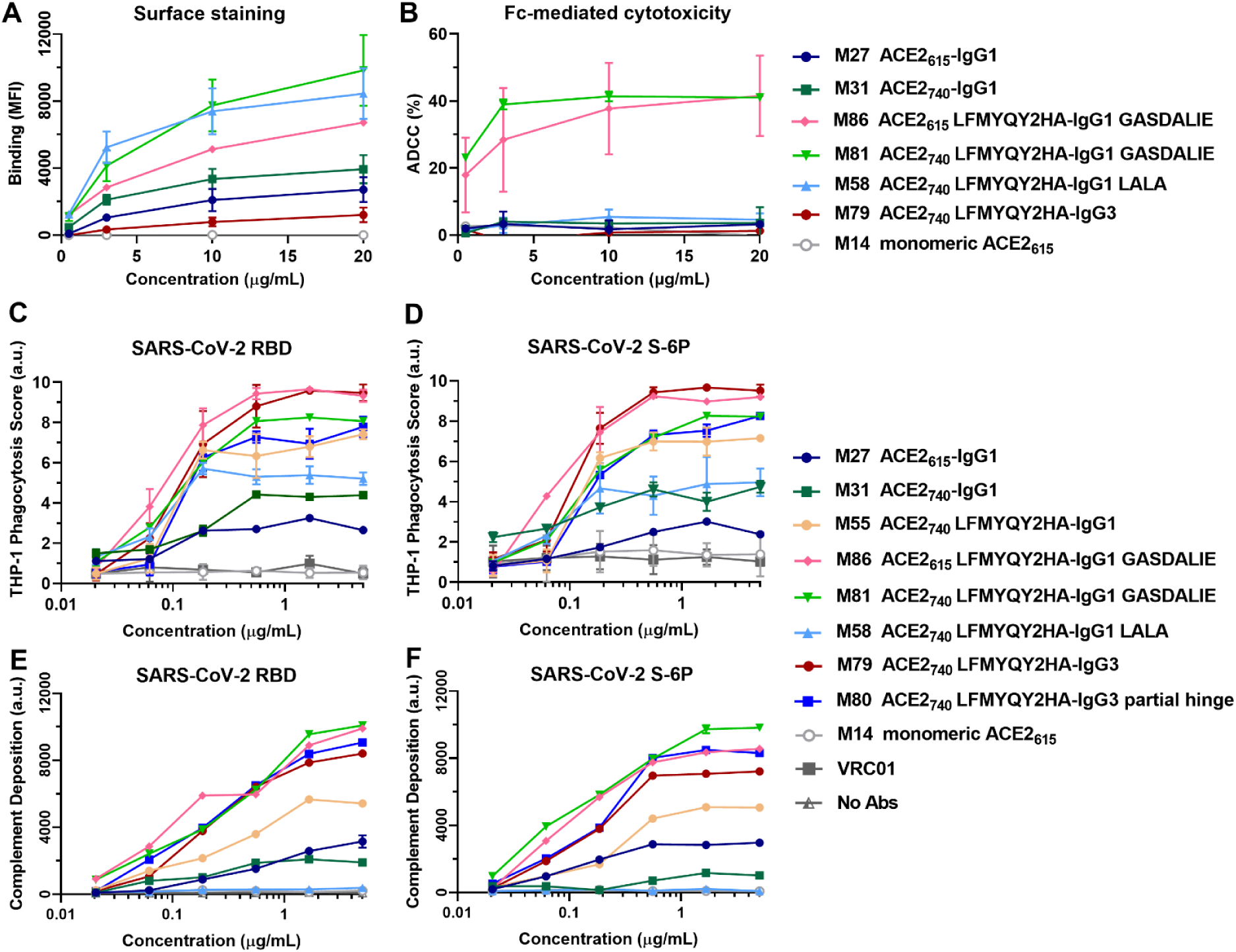
Engineered ACE2-Fcs mediate potent Fc-dependent cytotoxicity and phagocytosis *in vitro*. (**A**) Mean Fluorescence Intensity (MFI) of CEM.NKr cells expressing SARS-CoV-2 S (CEM.NKr-S) stained with indicated concentrations (0.5-20 μg/mL) of ACE2-Fcs or monomeric ACE2. The background MFI signal obtained on parental CEM.NKr CCR5+ cells was subtracted to the signal on CEM.NKr.Spike cells. (**B**) Percentage of ADCC in the presence of titrated amounts of ACE2-Fcs or monomeric ACE2 as in (A) using 1:1 ratio of parental CEM.NKr CCR5+ cells and CEM.NKr-S cells as targets when PBMCs from healthy donors were used as effector cells. (**C-D**) Fc-dependent cellular phagocytosis by THP-1 effector cells against the fluorescent microspheres (1 μm) coated with SARS-CoV-2 RBD (**C**) or S-6P (**D**) in the presence of varying concentrations (0.02-5 μg/mL) of ACE2-Fcs, monomeric ACE2 or VRC01. Data were the mean from at least 2 technical replicates. (**E-F**) Fc-mediated complement deposition. Multiplex assay microspheres coated with SARS-CoV-2 RBD (**E**) or S-6P (**F**) were incubated with 4-fold serial diluted ACE2-Fcs, monomeric ACE2 (0.02-5 μg/mL) or blank buffer control (No Abs) prior to incubation with guinea pig complement. Anti-guinea pig C3 IgG (conjugated with a red pigment) was used to detect the bound C3 on immune complexes. Data were the mean from two independent replicates.

Unlike ADCC, all ACE2-Fc variants showed dose-dependent ADCP activity against beads coated with both viral antigens (**Fig. 6 C-D**). M86 (ACE2_615_ LFMYQY2HA-Fc GASDALIE) and M79 (ACE2_740_ LFMYQY2HA -Fc in IgG3 backbone with full hinge) stimulated the highest phagocytic responses, followed by M81, a variant with longer ACE2 domain (ACE2_740_ LFMYQY2HA -Fc GASDALIE) and M80, ACE2_740_ LFMYQY2HA in IgG3 backbone but with a truncated hinge. This suggests that FcγR-Fc engagement of ACE2-Fc variants is partially impaired by increased flexibility of the CLD (M86 versus M81) or shorter hinge length (M79 versus M80). The latter observation is consistent with previously published data showing several HIV antibodies with extended hinges mediated enhanced ADCP against gp140-coated beads (*42*). Of note, residual ADCP activity for M58, the Fc-LALA variant, was observed against antigen-coated beads (**Fig. 6C-D**) and S-expressing CEM.NKr target cells (**Fig. S9**), respectively. This result agrees with reported data indicating that LALA mutations do not completely abrogate binding to FcγRII, a primary receptor involved in ADCP signaling (*61, 62*).

In the bead-based ADCD assay (**Fig. 6E-F**), Fc-mediated complement activation was affected by: 1) ACE2-Fc binding capacity to antigens coupled on the beads, 2) Fc subclass, and 3) ACE2-Fc concentrations. The LFMYQY2HA bearing M81 and M86 with GASDALIE Fc elicited the most potent deposition of complement C3, followed by the IgG3 subtypes M79 and M80. M55 with LFMYQY2HA but without GASDALIE mutation showed significantly weaker ADCD activity and its Fc LALA (M58) exhibited activity only slightly above baseline. The ACE2-Fc with no RBD binding enhancing mutations (M27 and M3) displayed only ∼30% ADCD efficiency as compared to M81 and M86. In summary, the ADCD efficiency of ACE-Fc was positively correlated with its binding affinity to the antigen-coated beads, as well as Fc composition, with a C1q recruitment efficiency rank of IgG1 GASDALIE-Fc > IgG3 > IgG1 >IgG1 LALA-Fc.

### Engineered ACE2-Fc with enhanced Fc effector functions protects or delays SARS-CoV-2 lethal infection in K18-hACE2 mice

We next tested if the engineered hACE2-Fc mediated protection when administered prophylactically or therapeutically in K18-hACE2 mice challenged with SARS-CoV-2-nLuc (WA/2020) and if *in vivo* efficacy was associated with Fc-mediated effector functions. K18-hACE2 mice were treated with vehicle (PBS or human IgG1 isotype control) or two engineered ACE2-Fcs, i.e. ACE2_740_ LFMYQY2HA-Fc GASDALIE (M81) or ACE2_740_ LFMYQY2HA –Fc LALA (M58), both intranasally (i.n., 12.5 mg/kg) and intraperitoneally (i.p., 6.25 mg/kg) 5 h before (prophylaxis) or 24 h after (therapeutic) i.n challenge with SARS-CoV-2-nLuc (1 x 10^5^ FFU) (**Fig. 7A**). Longitudinal non-invasive BLI and terminal imaging revealed that prophylactic administration of M81 efficiently inhibited SARS-CoV-2 replication in the lungs, nose and prevented neuro-invasion as compared to control mice (**Fig. 7B-D, S10A-B**). While control mice showed rapid weight loss and succumbed to infection at 6 dpi (**Fig. 7E-F**), three out of the four mice treated with M81 (Fc GASDALIE) did not experience any weight loss and were completely protected, and the fourth showed a significant delay in weight loss and survived until 11dpi. (**Fig. 7B and C**). Although all mice pre-treated with M58 (Fc LALA) showed delayed weight loss during initial phase of infection, only one out of four mice fully recovered (**Fig. 7B-F**). These results revealed a significant difference in the prophylactic outcome for mice pretreated with the GASDALIE variant, engineered to have enhanced affinity for Fcγ receptors and thus elevated Fc-effector activities including ADCC, ADCP and ADCD (**Fig. 8**), to those pretreated with the LALA variant which are capable of direct neutralization (**Fig. 6**) but impaired in Fc-effector functionality. Altogether these data demonstrated that both direct neutralization and Fc-effector activities of ACE2-Fc contributed to protection from infection and death of mice in the prophylactic setting (**Fig. 7B-C**).

**Fig. 7.**
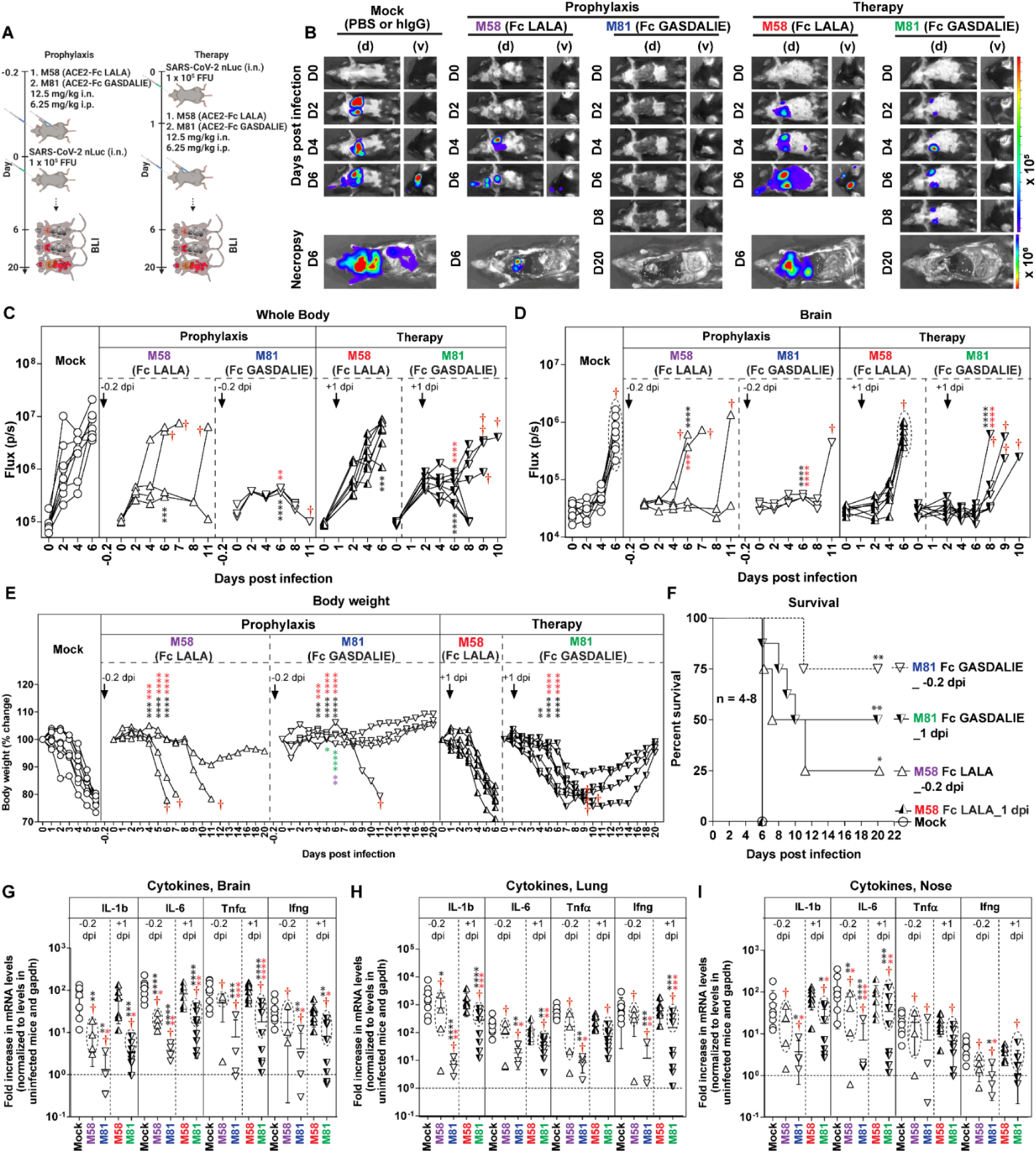
*In vivo* efficacy of Fc-null/enhancing ACE2-Fcs in prevention from lethal SARS-CoV-2 infection in K18 hACE2 transgenic mice. (**A**) A scheme showing the experimental design for testing the *in vivo* efficacy of M58 (ACE2_740_ LFMYQY2HA -Fc LALA) and M81 (ACE2_740_ LFMYQY2HA -Fc GASDALIE) delivered with a dose of 12.5 mg/kg body weight intranasally (i.n.) and 6.25 mg/kg body weight intraperitoneally (i.p.) 5 h before infection, (−0.2 dpi, prophylaxis) or 1 day after (+1 dpi, therapy) challenge of K18-hACE2 mice with 1 x 10^5^ FFU SARS-CoV-2-nLuc. PBS (n=4) or human IgG-treated (n=4) mice were used as controls. (**B**) Representative BLI images of SARS-CoV-2-nLuc-infected mice in ventral (v) and dorsal (d) positions. (**C-D**) Temporal quantification of the nLuc signal as flux (photons/sec) computed non-invasively. (**E**) Temporal changes in mouse body weight with initial body weight set to 100% for the experiments shown in (**A**). Mice that succumbed to infection (cohorts not 100% mortality) are denoted with red dagger. (**F**) Kaplan-Meier survival curves of mice (n = 4-8 per group) statistically compared by log-rank (Mantel-Cox) test for the experiments shown in (**A**). (**G-I**) Fold changes in cytokine mRNA expression in brain, lung and nasal cavity tissues. Data were normalized to Gapdh mRNA in the same sample and that in non-infected mice after necropsy. Cytokines in indicated tissues were determined when they succumbed to infection (dashed ellipse with red dagger) and at 20 dpi for surviving mice. Grouped data in (C-E), (G-I) were analyzed by 2-way ANOVA followed by Tukey’s multiple comparison tests. Statistical significance for group comparisons to control are shown in black, M58 (prophylaxis) in purple, M81 (prophylaxis) in blue, M58 (therapy) in red and M81 (therapy) in green. Non-significant comparison are not shown. ∗, *P* < 0.05; ∗∗, *P* < 0.01; ∗∗∗, *P* < 0.001; ∗∗∗∗, *P* < 0.0001; Mean values ± SD are depicted.

Next, we tested if M81 or M58 can resolve established infection when administered one day after SARS-CoV-2 challenge (**Fig. 7A-B**). While M58 (Fc LALA) completely failed to rescue SARS-CoV-2 infected mice, treatment with M81 (Fc GASDALIE) protected 50% of mice and significantly delayed weight loss and death of mice that succumbed to infection (**Fig. 7B**). The protection correlated with significantly reduced viral replication in the brain, lungs and nose as determined by imaging (nLuc activity in organs; flux) after terminal necropsy as well as by viral load analyses (**Fig. S10A-D**). Concomitant with the efficient virus clearance, substantial reductions in mRNA level of selected pro-inflammatory cytokines in target organs nose, lung and brain were observed in M81-treated mice compared to mock and M58-treated cohorts (**Fig. 7G-I**). Importantly, Fc-effector activities played a predominant role in clearing established infection as all mice treated with ACE2–Fc LALA were succumbed to infection. Thus, enhanced Fc effector functions recruited by M81 ACE2-Fc were critical for virologic control during therapy.

## DISCUSSION

ACE2-based therapies have proven effective at countering SARS-CoV-2 infection in humanized organoids (*22*), hamsters (*28*) and in a human clinical trial of a severely infected patient (*24*). These interventions include soluble monomeric ACE2 as well as dimeric ACE2 immunoglobulin-like molecules. As the primary host receptor, recombinant ACE2 derivatives are intrinsically broad neutralizers of ACE2-utilizing coronaviruses which include SARS-CoV-1, SARS-CoV-2, and CoVs found in bats with pandemic potential. Furthermore, human ACE2 decoys have an advantage over antibodies elicited by infection or vaccination as they are predisposed to block mutational escape of VOCs that emerge to increase infectivity by enhancing affinity to the ACE2 receptor. Here, we developed bifunctional ACE2-Fc variants that not only broadly neutralize SARS-CoV-2 VOCs but also engage host innate immune cells through the Fc to efficiently eliminate free virions or SARS-CoV-2-infected cells. We also show that both modes of action are required for ACE2-Fc to optimally prevent and control lethal SARS-CoV-2 infection in a K18-hACE2 mouse model.

Our ACE2-Fc variants’ design was guided by a structure-based approach to identify ACE2 mutations that enhance the affinity for the SARS-CoV-2 S glycoprotein. We identified several novel ACE2 mutations that facilitated RBD binding by up to ∼13 fold and improved neutralization potency and breadth against seven SARS-CoV-2 VOCs in PsV assays, including the prevalent Delta variant, B.1.617. Our lead variant, ACE2_740_ LFMYQY2HA–Fc GASDALIE, consists of three mutations, L79F, M82Y, and Q325Y, in the RBD-interacting region of ACE2; L79F and M82Y are in the Site II region of ACE2 and stabilize the mobile RBD ridge while Q325Y sits in the Site III region and introduces a new hydrogen bond to the binding interface. The ACE2 Site I helix has already been extensively used by others to generate several ACE2 mutants with higher affinity to the RBD (*25, 27–29*). Our additional mutations to Sites II and III could therefore potentially be combined with Site I mutations to make an even more potent ACE2 antiviral for use against the constantly evolving SARS-CoV-2 virus. As described by Higuchi, *et al* (*28*), the immunogenicity of engineered ACE2-Fcs, e.g., inducing adverse T cell activation and auto-antibodies that target the endogenous ACE2, need to be examined with caution.

We aimed at designing ACE2-Fc variants with eliminated ACE2 angiotensin enzymatic activity. By introducing two mutations (H374A and H378A) in the catalytic zinc binding site we generated variant with significant rearrangements to the substrate binding site that eliminated the unwanted proteolytic activity. The crystal structure of this engineered ACE2 in complex with SARS-CoV-2 RBD provided confirmation at the molecular level for this lack of activity. At present, it is unclear as to whether the enzymatic activity of ACE2 should be retained in ACE2 based therapeutics used to treat SARS-CoV-2 patients. Given that recombinant soluble ACE2 has been found to be safe and without obvious hemodynamic impact in healthy volunteers and that ACE2 activity has correlated with improved clinical outcome with regard to ACE-induced lung injury in ARDS patients (*63–65*), the enzyme activity of hrsACE2 is thought to be beneficial to SARS-CoV-1 and SARS-CoV-2 infected patients to alleviate severe lung injury when membrane-bound ACE2 has been stripped and downregulated by S binding (*6, 66*). On the other hand, a marked reduction of Ang II and an increase in Ang 1-7 were observed throughout the entire recombinant soluble ACE2 treatment period in one SARS-CoV-2 patient (*24*) which potentially increases the risk of hypotension. Presumably, repeated delivery of recombinant soluble ACE2 with enzyme activity may not only deregulate the RAS hormonal cascade but could also potentially downregulate the endogenous expression of surface ACE2. Therefore, until the risk to benefit ratio is thoroughly explored for the role of ACE2 activity in SARS-CoV-2 infection, inactive ACE2-derivatives serve as a safer alternative for therapeutic consideration.

Our PsV-challanged K18-hACE2 mouse model, adapted from a previously reported protocol (*56*), provided a safe and inexpensive platform for the dynamic *in vivo* efficacy assessment of SARS-CoV-2 antivirals, which can be widely used in BSL-2 laboratories. With this preclinical model, the mutational effects elicited by the S protein of SARS-CoV-2 VOCs can be rapidly evaluated, although the viral transduction is only limited to the nasal cavity and high PsV titers are required for BLI visualization. In this model our best performing variant ACE2_740_ LFMYQY2HA –Fc GASDALIE (i.n. 5µg), was able to prevent viral transduction as effectively as 25 µg of unmutated ACE2-Fc.

A few ACE2-Fc variants have been developed by others as therapeutic candidates that are also capable of viral neutralization, mostly by mechanisms involving direct competition for viral S binding to the host cell surface ACE2 (*23, 25-29, 47*). These molecules were developed to have increased affinity for RBD to enhance host cell ACE2 competition and neutralization. However, ACE2-Fcs are engineered to act as IgGs and are not only capable of interacting with viral antigen bivalently, and therefore with higher avidity, but also ‘profit’ from the Fc domain that can be recognized by effector cells in the host. Interestingly, our data indicate that ACE2-Fc with ‘wild-type’ Fc has only moderate Fc-effector activity *in vitro*. The only Fc-effector activities we detected for our ACE2_740_ LFMYQY2HA –Fc(wt) variant were moderate ADCP, and antibody complement C3 deposition. Interestingly, no ADCC of SARS-CoV-2 S-expressing T-lymphoid cells was detected for any of our optimized ACE2-Fc(wt) variants. Potent ADCC, enhanced ADCP, and complement activation were only detected when ACE2-Fcs were modified to include the well-known, low affinity Fcγ receptor enhancing GASDALIE mutations (*44*). Interestingly, ACE2_740_ LFMYQY2HA–Fc GASDALIE in an IgG1 backbone had ADCP activities comparable to ACE2_740_ LFMYQY2HA–Fc in an IgG3 backbone, pointing toward the possibility that like ADCP in HIV-1, IgG isotype and hinge length play a role in ADCP (*42, 43*).

Thus far, the *in vivo* protective potential of an ACE2-Fc therapeutic has been tested only once in a Syrian hamster model (*28*) which has several limitations due to its inability to fully recapitulate SARS-CoV-2 pathogenesis and severity. To better test our lead ACE2-Fc variant we utilized a well characterized K18-hACE2 mouse model (*67*). Due to the constitutive high endogenous human ACE2 expression, this model is highly susceptible to SARS-CoV-2 infection and the disease progression partially recapitulates the severe pathological features of SARS-CoV-2 infection in humans. The model has also been used extensively for evaluating contributions from direct neutralization and Fc-effector activities mediated by nAbs (*68*) and a non-neutralizing Ab (*69*). However, a high basal level of hACE2 on target cells in this model, particularly in the brain, poses a significant obstacle for soluble ACE2-based antivirals such as our engineered ACE2-Fc to surmount and achieve protection. Despite these limitations, we detected a strong benefit to the administration of ACE2_740_ LFMYQY2HA–Fc GASDALIE variant both prophylactically and therapeutically in K18-hACE2 mice. In both settings, ACE2-Fc treatments were associated with markedly improved *in vivo* efficacy, e.g., a reduction in virus-induced body weight loss, pro-inflammatory cytokine responses and mortality, particularly in the therapeutic context. Given the human Fc-mouse FcγR mismatch may compromise Fc-effector functionality of ACE-Fcs in K18-hACE2 mice, we expect a better therapeutic outcome in species matched systems, such as humanized-FcγR mice or clinical trials. Importantly, we did not observe any Fc-related pathogenic or disease-enhancing effects in ACE2-Fc treated mice, although recent studies have revealed a potential link between higher FcγRIII activation and disease severity with elevated afucosylated IgG levels in hospitalized SARS-CoV-2 patients (*70, 71*). Further studies are required to delineate the Fc-effector functions and Fc-FcγR-mediated pathways that confer the improved efficacy of Fc-engineered human IgGs and Fc-fusion molecules considering their immunomodulatory, inflammatory, and cytotoxic activity in other settings.

To summarize, our data confirm the utility of engineered ACE2-Fcs as valuable therapeutic agents capable of countering SARS-CoV-2 infection when administered prophylactically and therapeutically. Importantly, our data point toward a crucial role of Fc-effector activity in mechanism of anti-viral action of ACE2-Fc. While the engineered ACE2-Fc in wild-type IgG1 backbone showed moderate Fc-functions *in vitro*, the equivalent Fc-enhancing variants robustly stimulate Fc-effector responses and confer improved *in-vivo* protection. Altogether, as has been demonstrated for many nAbs, our findings strengthen the translational relevance of engineered ACE2-Fcs with improved Fc-effector functions as first-line antivirals for mild to moderate SARS-CoV-2 infection and highlights the importance of Fc-mediated effector functions in their mechanism of protection.

## MATERIALS AND METHODS

### Plasmids construction

The ACE2-RBD interfaces of two reported crystal structures (6M0J and 6VW1) were analyzed by PISA (*72*). To generate the expression plasmids of human ACE2 and human IgG1 fusions, the synthetic gene (GenBank BAJ21180.1, with original BamHI site destroyed) encoding the human ACE2 PD (residue 1-615) and the complete extracellular domain (residue 1-740, ECD) were fused to the human IgG1 Fc segment (residue D217-K443) or the codon optimized human IgG3 Fc (GenBank: AIC59039.1) with full hinge (residue E243-G520, R509H) or partial hinge (residue P286-G520, R509H), in which a BamHI site was inserted between the ACE2 and IgG Fc. The DNA chimera was then cloned into the pACP-tag (m)-2 vector (addgene# 101126) using NheI and NotI (NEB) as the restriction sites. All ACE2 mutations were introduced onto the ACE2-IgG backbone by a two-step mutagenesis protocol, described in (*73*). Likewise, to generate the engineered-Fc variants, gene segments encoding ACE2 PD, ECD or those with desired ACE2 mutations were fused to the codon-optimized synthetic IgG1 Fc (GenScript) in which GASDALIE or LALA mutations were incorporated. To generate SARS-CoV-2 RBD_wt_ (residue 319-541 or residue 319-537, for crystallization), RBD_B.1.1.7_ (residue 329-527, N501Y) and RBD_B.1.351_ (residue 329-527, K417N/E384K/N501Y), the respective codon optimized DNA segments fused with an N-terminal secretion peptide and a C-terminal 6xHis tag were cloned into the pACP-tag (m)-2 vector using either EcoRI/NotI for RBD_wt_ (319–541), RBD _B.1.1.7_ and RBD_B.1.351_ or BamHI/XhoI for RBD_wt_ (319–537) as restriction enzymes.

### Protein expression and purification

FreeStyle 293F cells (Thermo Fisher Scientific) were grown in FreeStyle 293F medium (Thermo Fisher Scientific) to a density of 1×10^6^ cells/mL at 37°C with 8% CO_2_ with 135 rpm agitation. For production of ACE2-Fc variants, cells were transfected with the corresponding plasmids (100ug/10^8^ cells) following the polyethylenimine (PEI) transfection protocol described in (*74*). One-week post-transfection, cells were pelleted and supernatant was clarified using a 0.22-µm filter and protein was purified using Protein A resin (Pierce), followed by size-exclusion chromatography (SEC) on Superose 6 10/300 column (Cytiva) equilibrated with 1x phosphate-buffered saline (PBS). Monomeric ACE2_wt_ and engineered ACE2_LFMYQY2HA_ plasmids encoding ACE2 (residue 1-615) with C-terminal HRV-3C-cleavable 8xHis tag (*45*) were transfected to FreeStyle 293F cells and the resulting protein was purified over Ni-NTA columns (Cytiva). His-tag removal was carried out by overnight HRV-3C (Sigma) digestion at 4°C and the cleaved protein was then purified on Ni-NTA before being subjected to SEC on Superose 6 10/300 column (Cytiva) equilibrated with PBS.

For recombinant expression of SARS-CoV-2 stabilized spikes ecto-domain(S-2P (*45*) and S-6P (*75*), gifted from Dr. Jason S. McLellan), RBD_wt_ (residue 319-541), RBD_B.1.1.7_, RBD_B.1.351_ and SARS-CoV RBD (residue 306-577, with C-terminal HRV3C-cleavable IgG1 Fc tag and 8xHis tag) (*45*), plasmids encoding the respective genes were transfected to 293F cells with the same protocol as described above. Supernatants were purified on either StrepTactin resin (IBA) for S-2P and S-6P or Ni-NTA columns for SARS-CoV RBD, SARS-CoV-2 RBD and its variants. S-2P, S-6P and SARS-CoV RBD were then incubated with HRV3C protease at 4 °C overnight and the mixtures were passed over a Ni-NTA column to remove the protease and cleaved tags. All viral proteins were further purified by SEC on either a Superose 6 10/300 or a HiLoad 16/600 Superdex 200 pg in PBS before being used for indirect ELISA and surface plasma resonance.

### Single-molecule mass photometry

The sample quality and molecular weight (M.W.) of the glycosylated monomeric ACE2_615_, ACE2-Fcs and SARS-CoV-2 S-6P were assessed by mass photometry (MP). Purified non-tagged ACE2_615_, ACE2-Fcs or S-6P were diluted to ∼50nM in PBS and MP data were acquired and analyzed using a OneMP mass photometer (Refeyn Ltd, Oxford, UK). The estimated M.W. (75 kD, 230 kD, 270 kD and 540 kD for ACE2_615_, ACE2_615_-Fc, ACE2_740_-Fc and SARS-CoV-2 S-6P, respectively) were used for A280-based concentration determination (corrected by extinction coefficients).

### ELISA

Binding capacity of the purified ACE2-Fcs to various viral antigens were measured by indirect ELISA, as described in (*76*). 96-well Nunc Maxisorp plates (Sigma) were coated with SARS-CoV-2 RBD_wt_ (residue 319-541) (50ng), RBD_B.1.351_ (50 ng), S-2P (75 ng), S_B.1.1.7_ (75 ng), S_B.1.351_ (75ng), S_P.1_ (75 ng), S_B.1.526_ (75 ng) and SARS-CoV RBD (50ng) per well in Tris-buffered saline (TBS) at 4 °C overnight. Plates were washed with TBS before blocking with TBS + 5% non-fat milk powder and 0.1% Tergitol (blocking buffer) at room temperature for 2 h. After 1x washing by TBS supplemented with 0.1% Tween 20 (TBST), serial dilutions of purified ACE2-Fcs (125, 62.5, 20, 10.0, 5.0, 2.5, 0.5 0.05 nM) were added and incubated at 4 °C overnight. Plates were washed three times and incubated with the goat anti-human-IgG Fc secondary antibody conjugated with alkaline phosphatase (AP, Southern Biotech) at a 1:1000 dilution in blocking buffer for 1 h at room temperature. Plates were washed three times and developed using the Blue Phos Microwell Phosphatase Substrate System (SeraCare). The reactions were stopped after 5 min incubation at room temperature by adding the equivalent volume of APstop Solution (SeraCare). The plates were then read at 620 nm and the optical density recorded by the SpectraMax Plus microplate reader (Molecular Devices). All binding events were measured in triplicate and each data set was normalized (OD_620_ at 125 nM as 100%) for cross-comparison. GraphPad Prism was used to display the mean and SEM for all groups and used to calculate the area under the curve (AUC) within the concentration range of 0.05-2.5 nM using 5% binding as baseline (**Fig. 2D** **& S2**).

### Surface Plasmon Resonance (SPR)

SPR measurements were done following carried out as described in (*68*). All assays were performed on a Biacore 3000 (Cytiva) at room temperature using 10 mM HEPES pH 7.5, 150 mM NaCl, 0.05% Tween 20 as running buffer. For the kinetic measurement of SARS-CoV-2 RBD_wt_, RBD_B.1.1.7_, RBD_B.1.351_ and SARS-CoV RBD binding to ACE2-Fc variants, ∼80-200 RU of ACE2-Fcs were immobilized on a Protein A chip (Cytiva) and 2-fold serial dilutions of the respective viral proteins were then injected as solute analytes with concentrations ranging from 6.25-200 nM (SARS-CoV-2 RBD_wt_ and SARS-CoV RBD) or 3.125-200 nM (RBD_B.1.1.7_ and RBD_B.1.351_). For kinetic measurement of the non-tagged SARS-CoV-2 S-6P binding to ACE2-Fcs, ∼60 RU of ACE2-Fcs were loaded on a Protein A chip before the serial injection of 2-fold titrated S-6P (3.125-50 nM). To assesFor monomeric ACE2_wt_ or ACE2_LFMYQY2HA_ binding to SARS-CoV-2 RBD-Fc, ∼120 RU of SARS-CoV-2 RBD_wt_ (residue 319-591) was immobilized on a Protein A chip and 2-fold serial dilutions of monomeric ACE2 or the variant were injected with concentrations ranging from 6.25-100 nM. For all kinetic assays, the sensor-chip was regenerated using 10mM Glycine pH 2.0 before the next cycle. Sensorgrams were corrected by subtraction of the corresponding blank channel as well as for the buffer background and kinetic constants were determined using a 1:1 Langmuir model with the BIAevaluation software (Cytiva), as shown in **Fig. 2E-F** and **S3**. The kinetic constants are summarized in **Table S1**. Goodness of fit of the curve was evaluated by the Chi^2^ of the fit with a value below 3 considered acceptable.

### Crystallization and structure determination

For crystallographic protein preparation, plasmids encoding ACE2 (residue 1-615, LFMYQY2HA, with C-terminal HRV3C-cleavable 8xHis tag) or SARS-CoV-2 RBD (residue 319-537) were transfected into Expi293F GnTI-Cells (Thermo Fisher Scientific) using PEI. The proteins were harvested and purified on Ni-NTA, the C-terminal 8xHis tag on ACE2 was removed by HRV3C digestion as above. The resulting SARS-CoV-2 RBD and the cleaved ACE2 were further purified by gel filtration on Superose 6 10/300 in PBS.

The purified non-tagged ACE2_615_(LFMYQY2HA) was mixed with excess RBD (molar ratio 1:5) and incubated on ice for 2h. The mixture was then deglycosylated by Endo H_f_ (NEB) in 1x PBS at room temperature overnight. Endo H_f_ was removed by repeated loading onto Amylose resin (NEB) and the crude ACE2-RBD mixture was further purified on a HiLoad 16/600 Superdex 200 which was pre-equilibrated in 10 mM Tris pH 8.0 and 100 mM ammonium acetate. The complex fractions were pooled and concentrated to ∼7.5 mg/mL for crystallization. Crystallization trials were performed using the vapor-diffusion hanging drop method with a 1:1.5 ratio of protein to well solution. Rod-shaped crystals were obtained in 0.2 M ammonium sulfate, 0.1 M MES pH 6.5, 20% (w/v) PEG 8000 after ∼3 weeks incubation at 21 °C. Crystals were snap-frozen in the crystallization condition supplemented with 20% MPD. X-ray diffraction data was collected at the SSRL beamline 9-2 and was processed with HKL3000 (*77*). The structure was solved by molecular replacement in PHASER from the CCP4 suite (*78*) using 6M0J (*37*) and 1R4L (*48*) as independent searching models for the RBD and ACE2 moiety respectively. Iterative cycles of model building and refinement were done in Coot (*79*) and Phenix (*80*). Data collection and refinement statistics are shown in **Table S2**. Structural analysis and Fig. generation were performed in PyMOL (*81*) and Chimera X (*82, 83*).

### ACE2 enzyme activity assay

Angiotensin converting activity was determined using the flurometirc ACE2 assay kit (BioVision). Briefly, the wtACE2-Fcs (M27 & M31) and H374A/H378A bearing M33 & M81 were diluted in assay buffer to the final concentrations of 1.56, 3.13, 6.25, 12.5, 25 and 50 nM and the reactions were set up with/without ACE2 inhibitors as described in the manufacturer protocol. The time-course measurements (Ex/Em=320/420 nm) were performed in the EnSpire multi-mode plate reader (Perkin Elmer). The initial linear regions in **Fig. S4D** were used to calculate the slopes d(RFU)/d(t) in given ACE2-Fc concentrations shown in **Fig. 3G**.

### Package of SARS-CoV-2 PsV

PsVs for *in vitro* neutralization assays, live cell imaging and *in vivo* efficacy studies were produced using the SARS-CoV-2 S-Pseudotyped Lentiviral Kit (NR-52948, BEI Resources) as described in (*50*). The resulting PsV lentiviral particles with SARS-CoV-2 S_wt_ expressed on the surface contained the reporter genes of synthetic firefly luciferase (Luc2) and synthetic *Zoanthus sp.* ZsGreen1. To generate PsV pseudotyped with spikes of different SARS-CoV-2 VOCs, Spike pseudotyping vector plasmids, including D614G (NR-53765, BEI Resources), P.1 (gift from Dr. Robert Petrovich, etc. from NIEHS), B.1.1.7, B1.351, B.1.429, P.1, B.1.526 and B.1.617.2. (InvivoGen), were used in lieu of the S_wt_ plasmid. 16-24h post seeding, 293T cells (Thermo Fisher Scientific) were co-transfected with respective spike plasmid or VSV G (positive control), lentiviral backbone and three helper plasmids encoding Gag, Tat1b and Rev1b (BEI Resources). At 72 h post transfection, the supernatant was harvested and clarified by 0.45-μm filters. To determine viral titers, hACE2-expressing 293T cells (gift from Dr. Allison Malloy, USUHS) were infected with serial PsV dilutions. 48-60 h post infection, luciferase signal was detected by the Bright-Go Luciferase Assay System (Promega) for titer estimations (*50*). PsV were concentrated by the homemade 4-fold lentivirus concentrator (protocol of MD Anderson) and stored at 4°C for short-term use or -20 °C for longer storage.

### *In vitro* neutralization assay

For *in vitro* neutralization assays, 50 μL serial dilutions of Synagis, monomeric ACE2 (M14), selected ACE2-Fcs (M27, M31, M81 and M86) (final concentration: 0.005-50 ng/μL) were pre-incubated with 50 μL SARS-CoV-2 spike PsV (∼10^6^ RLU/mL) of Wuhan-Hu-1 strain or seven VOCs in 96-well plates at 37 ℃ for 1 h. Subsequently, hACE2-expressing 293T cells (1.25 × 10^4^ cells/well) in 50 μL culture medium, were added and incubated at 37 °C for 48h. Microscopic live cell imaging for ZsGreen was performed by an All-in-One Fluorescence Microscope BZ-X (Keyence) (**Fig. 4C, S6-7**), and the luciferase signal was further measured by the Bright-Go Luciferase Assay System (Promega). Data analysis and normalization followed the protocol as described in (*84*).

### Antibody dependent cellular cytotoxicity (ADCC) assay

The assay was carried out as previously described (*57, 58*). Briefly, for evaluation of anti-SARS-CoV-2 ADCC activity, parental CEM.NKr CCR5+ cells were mixed at a 1:1 ratio with CEM.NKr-Spike cells. These cells were stained by AquaVivid (Thermo Fisher Scientific) for viability assessment and by a cell proliferation dye eFluor670 (Thermo Fisher Scientific) and subsequently used as target cells. Overnight rested PBMCs were stained with another cellular marker eFluor450 (Thermo Fisher Scientific) and used as effector cells. Stained effector and target cells were mixed at a 10:1 ratio in 96-well V-bottom plates. Titrated concentrations (0.5-20 µg/mL) of ACE2-Fc variants were added to the appropriate wells. The plates were subsequently centrifuged for 1 min at 300xg, and incubated at 37°C, 5% CO_2_ for 5 hours before being fixed in a 2% PBS-formaldehyde solution. Since CEM.NKr-Spike cells express GFP, ADCC activity was calculated using the formula: [(% of GFP+ cells in Targets plus Effectors)-(% of GFP+ cells in Targets plus Effectors plus antibody)]/(% of GFP+ cells in Targets) x 100 by gating on transduced live target cells. All samples were acquired on an LSRII cytometer (BD Biosciences) and data analysis performed using FlowJo v10 (Tree Star).

### Antibody dependent cellular phagocytosis (ADCP) assay

ADCP assays were carried out as previously described (*59*). Briefly, streptavidin-coated 1 µm fluorescent microspheres were coated with biotinylated SARS-CoV-2 S-6P or RBD_wt_(residue 319-541) overnight at 4 °C. Following washing, the beads were incubated with purified ACE2-Fcs at varied concentrations (0.02-5 µg/mL) for 3 h at 37 °C, and were analyzed in duplicate. For Monocyte ADCP, THP-1 cells were utilized as effectors cells. Cells were added to the bead/antibody mixture and incubated overnight to allow phagocytosis. Samples were then fixed and analyzed via flow cytometry to define the fraction and fluorescent intensity of cells that phagocytosed one or more beads.

To estimate ADCP efficiency for cellular elimination, CEM.NKr-Spike cells were used as target cells that were labelled with a cellular dye (cell proliferation dye eFluor450). THP-1 cells were used as effector cells and were stained with another cellular dye (cell proliferation dye eFluor670). Stained target and effector cells were mixed at a 5:1 ratio in 96-well plates. Titrated concentrations (0.78-50 µg/mL) of ACE2-Fc variants were added to the appropriate wells. After an overnight incubation at 37 °C and 5% CO_2_, cells were fixed with a 2% PBS-formaldehyde solution. Antibody-dependent cellular phagocytosis was determined by flow cytometry, gating on THP-1 cells that were triple-positive for GFP, efluor450 and efluor670 cellular dyes. All samples were acquired on an LSRII cytometer (BD Biosciences) and data analysis performed using FlowJo v10 (Tree Star).

### Antibody-dependent complement deposition (ADCD) assay

As described in (*59*), the selected ACE2-Fc variants at varied concentrations (0.02-5 µg/mL) were incubated with multiplex assay microspheres coated with SARS-CoV-2 RBD or S-6P for 2hr at RT. Lyophilized guinea pig complement was resuspended according to manufacturer’s instructions (Cedarlane), and 2 μL per well was added in veronal buffer with 0.1% gelatin (Boston BioProducts). After washing, the mixtures of ACE2-Fc/microspheres were incubated with guinea pig complement serum at RT with shaking for 1 h. Samples were washed, sonicated, and incubated with goat anti-guinea pig C3 antibody conjugated with biotin (Immunology Consultants Laboratory) at RT for 1 h followed by incubation with streptavidin R-Phycoerythrin (PE, Agilent Technologie) at RT for 30min. After a final wash and sonication, samples were resuspended in Luminex sheath fluid and complement deposition was determined on a MAGPIX (Luminex Corp) instrument to define the median fluorescence intensity (MFI) of PE from two independent replicates. Assays performed without ACE2-Fc and without complement serum were used as negative controls.

### *In vivo* efficacy of ACE2-Fcs in K18-hACE2 mice challenged with SARS-CoV-2 PsVs

K18-hACE2 transgenic mice were purchased from The Jackson Laboratory. All mice were maintained under a specific pathogen-free (SPF) condition at the National Institute of Health Animal Facility. All animal experiments were performed according to Institute of Laboratory Animal Resources guidelines and the protocol was approved by the National Cancer Institute Animal Care and Use Committee.

For *in vivo* PsV-based inhibition assays, 6-8-week-old K18-hACE2 mice were intranasally (i.n) treated with Synagis (control IgG, 25 µg), M27 or M81 (5 or 25 µg) one hour before challenge by SARS-CoV-2 PsV_D614G_ or PsV_B.1.617.2_ (i.n., ∼10^8^ RLU). Dynamic luciferase signal was acquired 4, 8, and 12 dpi by IVIS® Spectrum In Vivo Imaging System (PerkinElmer) 10 min after i.n delivery of 200 μg D-luciferin (LUCK, GoldBio). Tissues (nasal cavity, trachea and lung) were collected 13 dpi and stored in -80 °C before processing.

For PK studies, C57BL/6J mice were intravenously (i.v.) injected with 100 μg (5 mg/kg) of two engineered ACE2-Fc M81 or M86. Before and after injection, serum samples were collected at 0 min, 10min, 1 h, 6 h, 24 h and 48 h and the ACE2-Fc serum concentration was estimated by indirect ELISA in which SARS-CoV-2 RBD_wt_ (200 ng/well) were used as capturing molecule and the goat-anti-human IgG conjugated with AP (1:1000 dilution) were used as secondary antibody. The alanine transaminase (ALT) and aspartate transaminase (AST) concentrations in sera before and 48 h after ACE2-Fc injection were assessed using commercial ALT and AST assay kits (Catachem) and monitored at 340 nm for 15 min with a microplate reader (BioAssay Systems).

For the quantitative real-time PCR of tissues from PsV-challenged mice, the tissues were lysed in TrizolTM Reagent (Invitrogen), and total RNA was extracted by phenol/chloroform. cDNA was synthesized from 1 μg total RNA using qSript cDNA SuperMix (Quantabio). The primer sequences applied in this study are listed in the **Table S3**. The relative level of each mRNA was calculated as fold change compared with control groups after normalizing with *Gapdh*.

### *In vivo* efficacy of ACE2-Fcs in K18-hACE2 mice challenged with SARS-CoV-2 nLuc

All experiments were approved by the Institutional Animal Care and Use Committees (IACUC) of and Institutional Biosafety Committee of Yale University (IBSCYU). All the animals were housed under specific pathogen-free conditions in the facilities provided and supported by Yale Animal Resources Center (YARC). hACE2 transgenic B6 mice (heterozygous) were obtained from Jackson Laboratory. 6–8-week-old male and female mice were used for all the experiments. The heterozygous mice were crossed and genotyped to select heterozygous mice for experiments by using the primer sets recommended by Jackson Laboratory.

For *in vivo* efficacy studies, 6 to 8 weeks old male and female mice were challenged i.n. with 1 x 10^5^ FFU SARS-CoV-2-nLuc WA/2020 in 25-30 µL volume under anesthesia (0.5 - 5 % isoflurane) delivered using precision Dräger vaporizer with oxygen flow rate of 1 L/min). For prophylaxis, purified ACE2-Fc proteins were administered i.n. at 12.5 mg/kg or 6.25 mg/kg for intraperitoneally (i.p.) injection, 5 h prior to infection of K18-hACE2 mice. For therapy, the same amounts (i.n and i.p) were administered 1 dpi. The starting body weight was set to 100 %. For survival experiments, mice were monitored every 6-12 h starting six days after virus administration. Lethargic and moribund mice or mice that had lost more than 20 % of their body weight, were sacrificed and considered to have succumbed to infection for Kaplan-Meier survival plots.

### Bioluminescence Imaging (BLI) of SARS-CoV-2 infection

All standard operating procedures and protocols for IVIS imaging of SARS-CoV-2 infected animals under ABSL-3 conditions were approved by IACUC, IBSCYU and YARC. All the imaging was carried out using IVIS Spectrum® (PerkinElmer) in XIC-3 animal isolation chamber (PerkinElmer) that provided biological isolation of anesthetized mice or individual organs during the imaging procedure. All mice were anesthetized via isoflurane inhalation (3 - 5 % isoflurane, oxygen flow rate of 1.5 L/min) prior and during BLI using the XGI-8 Gas Anesthesia System. Prior to imaging, 100 µL of nanoluciferase substrate, furimazine (NanoGlo^TM^, Promega, Madison, WI) diluted 1:40 in endotoxin-free PBS was retro-orbitally administered to mice under anesthesia. The mice were then placed into XIC-3 animal isolation chamber (PerkinElmer) pre-saturated with isoflurane and oxygen mix. The mice were imaged in both dorsal and ventral position on indicated dpi. The animals were then imaged again after euthanasia and necropsy by supplementing additional 200 µL of substrate on to exposed intact organs. Infected areas were identified by carrying out whole-body imaging after necropsy and were isolated, washed in PBS to remove residual blood and placed onto a clear plastic plate. Additional droplets of furimazine in PBS (1:40) were added to organs and soaked in substrate for 1-2 min before BLI.

Images were acquired and analyzed with Living Image v4.7.3 *in vivo* software package (Perkin Elmer Inc). Image acquisition exposures were set to auto, with imaging parameter preferences set in order of exposure time, binning, and f/stop, respectively. Images were acquired with luminescent f/stop of 2, photographic f/stop of 8 with binning set to medium. Comparative images were compiled and batch-processed using the image browser with collective luminescent scales. Photon flux was measured as luminescent radiance (p/sec/cm2/sr). Luminescent signals were regarded as background when minimum threshold setting resulted in displayed radiance above non-tissue-containing or known uninfected regions.

### Measurement of viral burden

Indicated organs (nasal cavity, brain and lungs) from infected or uninfected mice were collected, weighed, and homogenized in 1 mL of serum-free RPMI media containing penicillin-streptomycin and 1.5 mm Zirconium beads with a BeadBug 6 homogenizer (Benchmark Scientific, T Equipment Inc). Viral titers were measured using two highly correlative methods. First, the total RNA was extracted from homogenized tissues using RNeasy plus Mini kit (Qiagen), reverse transcribed with iScript advanced cDNA kit (Bio-Rad) followed by a SYBR Green Real-time PCR assay for determining copies of SARS-CoV-2 N gene RNA using primers are listed in **Table S3**.

Second, we used nanoluciferase activity as an efficient surrogate for a plaque assay. Dilutions from infected cell homogenates were applied on Vero E6 monolayer. 24 hour post infection, infected Vero E6 cells were washed with PBS, lysed with Passive lysis buffer and transferred into a 96-well solid white plate (Costar Inc) and nanoluciferase activity was measured using Tristar multiwell Luminometer (Berthold Technology) for 2.5 seconds by adding 20 µl of Nano-Glo® substrate in nanoluc assay buffer (Promega Inc). An uninfected monolayer of Vero E6 cells treated identically served as controls for determining background and obtain normalized relative light units. The data were processed and plotted using GraphPad Prism 8 v8.4.3.

### Analyses of signature inflammatory cytokines mRNA expression

Brain, lung and nose samples were collected from mice at the time of necropsy. Total RNA was extracted using RNeasy plus Mini kit (Qiagen), reverse transcribed with iScript advanced cDNA kit (Bio-Rad) followed by a SYBR Green Real-time PCR assay for determining the relative expression of selected inflammatory cytokines, i.e. *Il6, Ccl2, Cxcl10 and Ifng*, using primers listed in **Table S3**. The reaction plate was analyzed using CFX96 touch real time PCR detection system. The relative cytokine mRNA levels were calculated with the formula ΔC_t_(target gene) = C_t_(target gene)-C_t_(*Gapdh*). The fold increase was determined using 2-^ΔΔCt^ method comparing treated mice to uninfected controls.

### Quantification and Statistical Analysis

Data were analyzed and plotted using GraphPad Prism software (La Jolla). Statistical significance for pairwise comparisons were derived by applying non-parametric Mann-Whitney test (two-tailed). To obtain statistical significance for survival curves, grouped data were compared by log-rank (Mantel-Cox) test. To obtain statistical significance for grouped data we employed 2-way ANOVA followed by Tukey’s multiple comparison tests. *P* values lower than 0.05 were considered statistically significant. *P* values were indicated as ∗, *P* < 0.05; ∗∗, *P* < 0.01; ∗∗∗, *P* < 0.001; ∗∗∗∗, *P* < 0.0001.

## Disclaimer

The views expressed in this presentation are those of the authors and do not reflect the official policy or position of the Uniformed Services University, the U.S. Army, the Department of Defense, the National Institutes of Health, Department of Health and Human Services or the U.S. Government, nor does mention of trade names, commercial products, or organizations imply endorsement by the U.S. Government.

## Author contributions

Y.C., L.S., I.U., P.D.U. & M.P. conceptualized this study, design the experiments, analyzed data, generate figures and wrote the manuscript; Y.C. & M.P. designed the ACE2-Fc variants; Y.C., S.G., R.S., D.W. & D.N.N. produced, purified and characterized the proteins; Y.C. & S.M. performed SPR kinetics; Y.C. & W.D.T. solved and analyzed the crystal structure; L.S. & S.D. generated PsVs and performed neutralization assays; L.S. performed lived cell imaging; L.S., Y.L. & Y.C. designed, optimized and carried out *in-vivo* inhibition studies on PsV-challenged mice; G.B.B., S.P.A., A.P.H. & L.M. performed *in-vitro* ADCC, ADCP and ADCD assays; I.U. & P.D.U. designed, optimized and performed the in-vivo efficacy studies on SARS-CoV-2-nLuc infected mice; I.U., P.D.U. & L.S. analyzed the mice tissues, quantified viral loads and cytokines; R.S., W.D.T., A.F., G.B.B., S.P.A., D.N.N, M.E.A. and F.J.G. critically reviewed, edited and commented on the manuscript; M.P., F.J.G., P.D.U., W.M., A.F., M.E.A., G.P. & P.K. funded the work; Every author has read, edited and approved the final manuscript.

## Conflict of Interests

The authors declare that they have no competing interests.

## Data and materials availability

All data needed to evaluate the conclusions in the paper are present in the paper and/or the Supplementary Materials.

## Acknowledgements

We thank Dr. Jason S. McLellan from University of Texas, Austin for sharing the expression plasmids of SARS-CoV-2 S-2P, S-6P, SARS-CoV-1 RBD-Fc and monomeric ACE2. We thank BEI Resources for sharing the SARS-CoV-2 related reagents, including pseudotyped lentiviral kits (NR-53816 and NR-53817), recombinant S_B.1.1.7_ (NR-55311), S_B.1.351_ (NR-55311), S_B.1526_ (NR-55438), S_P.1_ (NR-55307), etc. We thank Dr. Kathleen Pratt and Dr. Pooja Vir from Department of Medicine, USUHS for providing laboratory resources and technical supports to this project. We thank Dr. Allison Malloy and Dr. Zhongyan Lu from Department of Pediatric, Uniformed Services University of the Health Science (USUHS) for sharing the hACE2-expressing HEK293T cell line. We thank Dr. Robert Petrovich and Dr. Negin P. Martin from NIEHS for sharing the S_P.1_ plasmid for PsV production.

## Funding

This work was supported by USUHS intramural funds to MP, in part by funding to F.J.G. from the Intramural Research Program, National Institutes of Health, National Cancer Institute, Center for Cancer Research, and in part by NIH grant R01AI163395 to W.M. A CIHR operating Pandemic and Health Emergencies Research grant #177958 and an Exceptional Fund COVID-19 from the Canada Foundation for Innovation (CFI) #41027 to A.F. A.F. is the recipient of Canada Research Chair on Retroviral Entry no. RCHS0235 950-232424. G.B.B. is the recipient of a FRQS PhD fellowship and S.P.A is the recipient of a CIHR PhD fellowship. Use of the Stanford Synchrotron Radiation Lightsource, SLAC National Accelerator Laboratory, is supported by the U.S. Department of Energy, Office of Science, Office of Basic Energy Sciences under Contract No. DE-AC02-76SF00515. The SSRL Structural Molecular Biology Program is supported by the DOE Office of Biological and Environmental Research, and by the National Institutes of Health, National Institute of General Medical Sciences.

## Notes

### Competing Interest Statement

The authors have declared no competing interest.

