## Supplementary Materials for "Engineered ACE2-Fc counters murine lethal SARS-CoV-2 infection through direct neutralization and Fc-effector activities"

**This PDF files includes:**

Table. S1-S3
Fig. S1 to S10
References (#1 to #6)

### Table S1. Summary of SPR kinetic constants

| Protein ID | Immobilized ligand | Flow antigen | Fitting mode | $k_{on}$ ×$10^4$ ($M^{-1}$·$s^{-1}$) | $k_{off}$ ×$10^{-3}$ ($s^{-1}$) | $K_D$ (nM) | $Chi^2$ value |
|---|---|---|---|---|---|---|---|
| **ACE2 (18-615)-Fc** | | | | | | | |
| M27 | Wild-type | SARS-CoV-2 RBD | 1:1 | 5.59 | 1.45 | 26.0 | 0.348 |
| | | RBD (B.1.1.7) | 1:1 | 16.7 | 4.14 | 24.8 | 0.67 |
| | | RBD (B.1351) | 1:1 | 32.3 | 3.73 | 11.6 | 0.66 |
| | | SARS-CoV-1 RBD | 1:1 | 45.7 | 57.5 | 126 | 1.61 |
| | | SARS-CoV-2 S-6P | 1:1 | 30.4 | 0.56 | 1.82 | 2.74 |
| M33 | H374A+H378A | SARS-CoV-2 RBD | 1:1 | 5.91 | 1.63 | 27.6 | 0.478 |
| | | RBD (B.1.1.7) | 1:1 | 24.9 | 5.86 | 23.60 | 1.43 |
| | | RBD (B.1351) | 1:1 | 46.5 | 3.58 | 7.70 | 0.26 |
| | | SARS-CoV-1 RBD | 1:1 | 48.3 | 56.0 | 116 | 3.05 |
| | | SARS-CoV-2 S-6P | 1:1 | 30.4 | 0.57 | 1.88 | 1.83 |
| M38 | L79F+M82Y | SARS-CoV-2 RBD | 1:1 | 63.7 | 2.98 | 4.67 | 0.725 |
| | | RBD (B.1.1.7) | 1:1 | 64.6 | 5.19 | 8.02 | 0.85 |
| | | RBD (B.1351) | 1:1 | 114 | 4.35 | 3.82 | 0.47 |
| | | SARS-CoV-1 RBD | 1:1 | 199 | 19.0 | 9.55 | 4.12 |
| | | SARS-CoV-2 S-6P | 1:1 | 33.7 | 0.198 | 0.587 | 2.33 |
| M39 | F28S+K31R | SARS-CoV-2 RBD | 1:1 | 4.27 | 2.72 | 63.61 | 1.45 |
| | | RBD (B.1.1.7) | 1:1 | 11.3 | 10.1 | 89.8 | 0.96 |
| | | RBD (B.1351) | 1:1 | 14.4 | 10.8 | 75.5 | 1.37 |
| | | SARS-CoV-1 RBD | 1:1 | N.D. | | | |
| | | SARS-CoV-2 S-6P | 1:1 | N.D. | | | |
| M40 | L45D | SARS-CoV-2 RBD | 1:1 | 49.0 | 9.63 | 19.66 | 1.01 |
| | | RBD (B.1.1.7) | 1:1 | 15.2 | 5.25 | 34.61 | 0.59 |
| | | RBD (B.1351) | 1:1 | 25.6 | 7.26 | 28.39 | 1.01 |
| | | SARS-CoV-1 RBD | 1:1 | N.D. | | | |
| | | SARS-CoV-2 S-6P | 1:1 | 19.5 | 0.34 | 1.76 | 0.73 |
| M41 | Q325Y | SARS-CoV-2 RBD | 1:1 | 42.5 | 3.88 | 9.14 | 0.842 |
| | | RBD (B.1.1.7) | 1:1 | 37.6 | 4.45 | 11.82 | 0.83 |
| | | RBD (B.1351) | 1:1 | 15.8 | 1.83 | 7.95 | 2.03 |
| | | SARS-CoV-1 RBD | 1:1 | 82.4 | 70.9 | 86 | 0.226 |
| | | SARS-CoV-2 S-6P | 1:1 | 19.6 | 0.42 | 2.14 | 1.56 |
| M86 | L79F+M82Y+Q325 | SARS-CoV-2 RBD | 1:1 | 102 | 3.64 | 3.58 | 0.596 |

| | | | | | | | |
|---|---|---|---|---|---|---|---|
| | Y+H374A+H378A LFMYQY2HA (GASDALIE-Fc) | RBD (B.1.1.7) | 1:1 | 105 | 5.04 | 4.78 | 0.56 |
| | | RBD (B.1351) | 1:1 | 110 | 1.75 | 1.59 | 0.53 |
| | | SARS-CoV-1 RBD | 1:1 | 143 | 18.5 | 12.9 | 0.0769 |
| | | SARS-CoV-2 S-6P | 1:1 | 37.0 | 0.197 | 0.533 | 2.39 |
| **ACE2 (18-740)-Fc** | | | | | | | |
| M31 | wild type | SARS-CoV-2 RBD | 1:1 | 43.3 | 6.0 | 13.9 | 1.08 |
| | | RBD (B.1.1.7) | 1:1 | 19.9 | 3.61 | 18.1 | 0.49 |
| | | RBD (B.1351) | 1:1 | 33.0 | 2.77 | 8.38 | 0.29 |
| | | SARS-CoV-1 RBD | 1:1 | 72.2 | 48.0 | 66.5 | 0.346 |
| | | SARS-CoV-2 S-6P | 1:1 | 12.9 | 0.13 | 1.02 | 1.23 |
| M58 | L79F+M82Y+Q325Y+ H374A+H378A LFMYQY2HA (LALA-Fc) | SARS-CoV-2 RBD | 1:1 | 84.6 | 2.83 | 3.35 | 1.26 |
| | | RBD (B.1.1.7) | 1:1 | 11.34 | 4.21 | 3.72 | 1.02 |
| | | RBD (B.1351) | 1:1 | 21.64 | 2.88 | 1.33 | 0.36 |
| | | SARS-CoV-1 RBD | 1:1 | 96.4 | 24.9 | 25.8 | 0.343 |
| | | SARS-CoV-2 S-6P | 1:1 | 40.9 | 0.365 | 0.891 | 1.51 |
| M79 | L79F+M82Y+Q325Y+ H374A+H378A LFMYQY2HA (IgG3-Fc) | SARS-CoV-2 RBD | 1:1 | 59.6 | 2.96 | 4.98 | 1.82 |
| | | RBD (B.1.1.7) | 1:1 | 38.9 | 6.21 | 16.0 | 1.01 |
| | | RBD (B.1351) | 1:1 | 71.0 | 3.39 | 4.77 | 0.37 |
| | | SARS-CoV-1 RBD | 1:1 | 107 | 23.6 | 22.1 | 0.406 |
| | | SARS-CoV-2 S-6P | 1:1 | 49.9 | 0.347 | 0.69 | 1.45 |
| M81 | L79F+M82Y+Q325 Y+H374A+H378A LFMYQY2HA (GASDALIE-Fc) | SARS-CoV-2 RBD | 1:1 | 81.8 | 2.24 | 2.74 | 1.51 |
| | | RBD (B.1.1.7) | 1:1 | 125 | 3.62 | 2.89 | 12.51 |
| | | RBD (B.1351) | 1:1 | 174 | 1.51 | 0.87 | 17.39 |
| | | SARS-CoV-1 RBD | 1:1 | 111 | 19.3 | 17.4 | 0.504 |
| | | SARS-CoV-2 S-6P | 1:1 | 43.4 | 0.33 | 0.76 | 0.964 |
| | SARS-CoV-2 S RBD (319-591)-Fc | Monomeric $ACE2_{615}$-wild type | 1:1 | 11.1 | 9.73 | 87.6 | 0.66 |
| | | Monomeric $ACE2_{615}$-LFMYQY2HA | 1:1 | 48.7 | 5.82 | 11.9 | 0.19 |

**Table S2. Crystallographic data collection and refinement statistics.**

| | $ACE2_{615}$ (LFMYQY2HA)-RBD |
|---|---|
| **Data collection** | |
| Wavelength, Å | 0.979 |
| Resolution range, Å | 47.51 - 3.54 (3.667 - 3.54) |
| Space group | $P2_1$ |
| Unit cell parameter | |
| a, b, c, Å | 132.6, 136.3, 132.7 |
| α, β, γ, ° | 90.0, 92.5, 90.0 |
| Redundancy | 22.1 (3.0) |
| Completeness, % | 96.98 (91.52) |
| Mean I/sigma(I) | 4.71 (1.23) |
| $R_{merge}$[a] | 0.151 (0.804) |
| $R_{pim}$[b] | 0.140 (0.896) |
| $CC_{1/2}$[c] | 0.899 (0.447) |
| Wilson $B_{factor}$, (1/Å$^2$)[d] | 108.39 |
| **Refinement** | |
| $R_{work}$[e] | 0.245 (0.362) |
| $R_{free}$[f] | 0.292 (0.411) |
| Resolution, Å | 47.51 - 3.54 |
| # of non-hydrogen atoms | |
| proteins | 25351 |
| water | 3 |
| Overall $B_{factor}$, (Å$^2$) | |
| proteins | 122.75 |
| ligands | 126.05 |
| water | 37.15 |
| RMS (bond lengths), Å | 0.004 |
| RMS (bond angles), ° | 0.69 |
| Ramachandran[g] | |
| Favored, % | 96.35 |
| Allowed, % | 3.61 |
| Outliers, % | 0.03 |
| PDB ID | 7RPV |

Statistics for the highest-resolution shell are shown in parentheses.

[a]$R_{merge} = \sum | I - \langle I \rangle | / \sum I$, where $I$ is the observed intensity and $\langle I \rangle$ is the average intensity obtained from multiple observations of symmetry-related reflections after rejections

[b]$R_{pim}$ = as defined in (*1*).

[c]$CC_{1/2}$ = as defined by Karplus and Diederichs (*2*)

[d]Wilson $B_{factor}$ as calculated in (*3*)

[e]$R = \sum \| F_o | - | F_c \| / \sum | F_o |$, where $F_o$ and $F_c$ are the observed and calculated structure factors, respectively.

[f]$R_{free}$ = as defined by Brünger (*4*)
[g]Calculated with MolProbity (*5*).

**Table S3. Sequences of the real time PCR primers**

| Target gene | Forward | Reverse |
|---|---|---|
| PsV *ZsGreen* | GACAGATAACTGGGAGCCATCC | CGGCATCTTTCTTGGCACAGAC |
| SARS-CoV-2 *N* | ATGCTGCAATCGTGCTACAA | GACTGCCGCCTCTGCTC |
| Mouse *Il1b* | GCCACCTTTTGACAGTGATGAG | CTCCTCTTCGCACTTCTGCTC |
| Mouse *Il6* | TAGTCCTTCCTACCCCAATTTCC | GACAGCCCAGGTCAAAGGTT |
| Mouse *Tnfa* | CCACCACGCTCTTCTGTCTAC | AGGGTCTGGGCCATAGAACT |
| Mouse *Ifng* | ATGAACGCTACACACTGCATC | CCATCCTTTTGCCAGTTCCTC |
| Mouse *Actin* | GGCTGTATTCCCCTCCATCG | CCAGTTGGTAACAATGCCATGT |

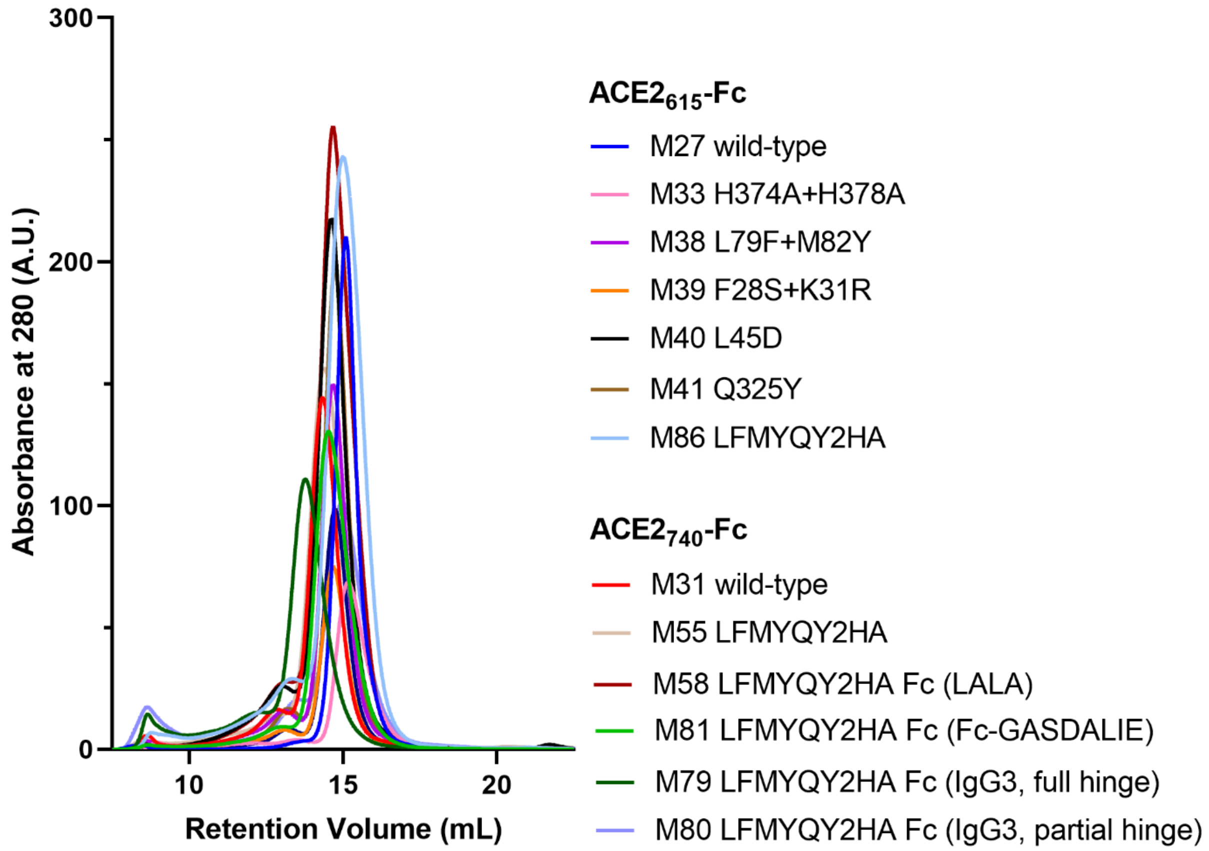

**Fig. S1. Size exclusion chromatographic (SEC) profiles of purified ACE2-Fc variants.**

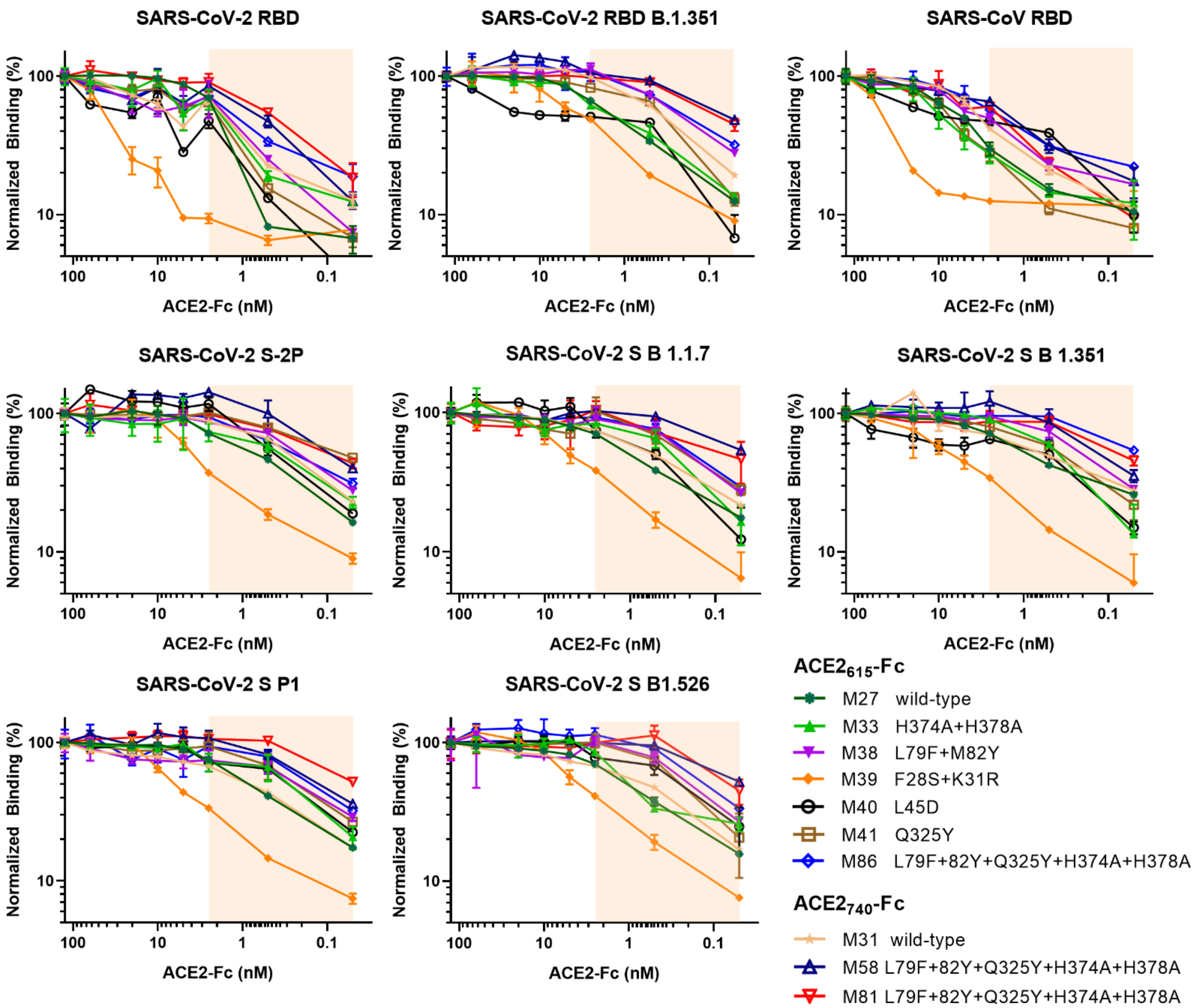

**Fig. S2. ELISA binding of ACE2-Fc variants to SARS-CoV-2 and SARS-CoV-1 antigens. Related to Fig. 2D.** Serial dilutions (0.05-125 nM) of purified ACE2-Fcs were applied to each well pre-coated with 50 ng of SARS-CoV-2 RBD, RBD$_{B1.351}$, SARS-CoV-1 RBD or 75 ng of SARS-CoV-2 S-2P, S$_{B.1.1.7}$, S$_{B.1.351}$, SP.1, S$_{B.1.526.}$ AUCs in the unsaturated region (0.05-2.50 nM, shaded in wheat) were calculated, normalized (binding at 125nM set as 100%) and plotted as heat-map shown in **Fig. 2D**.

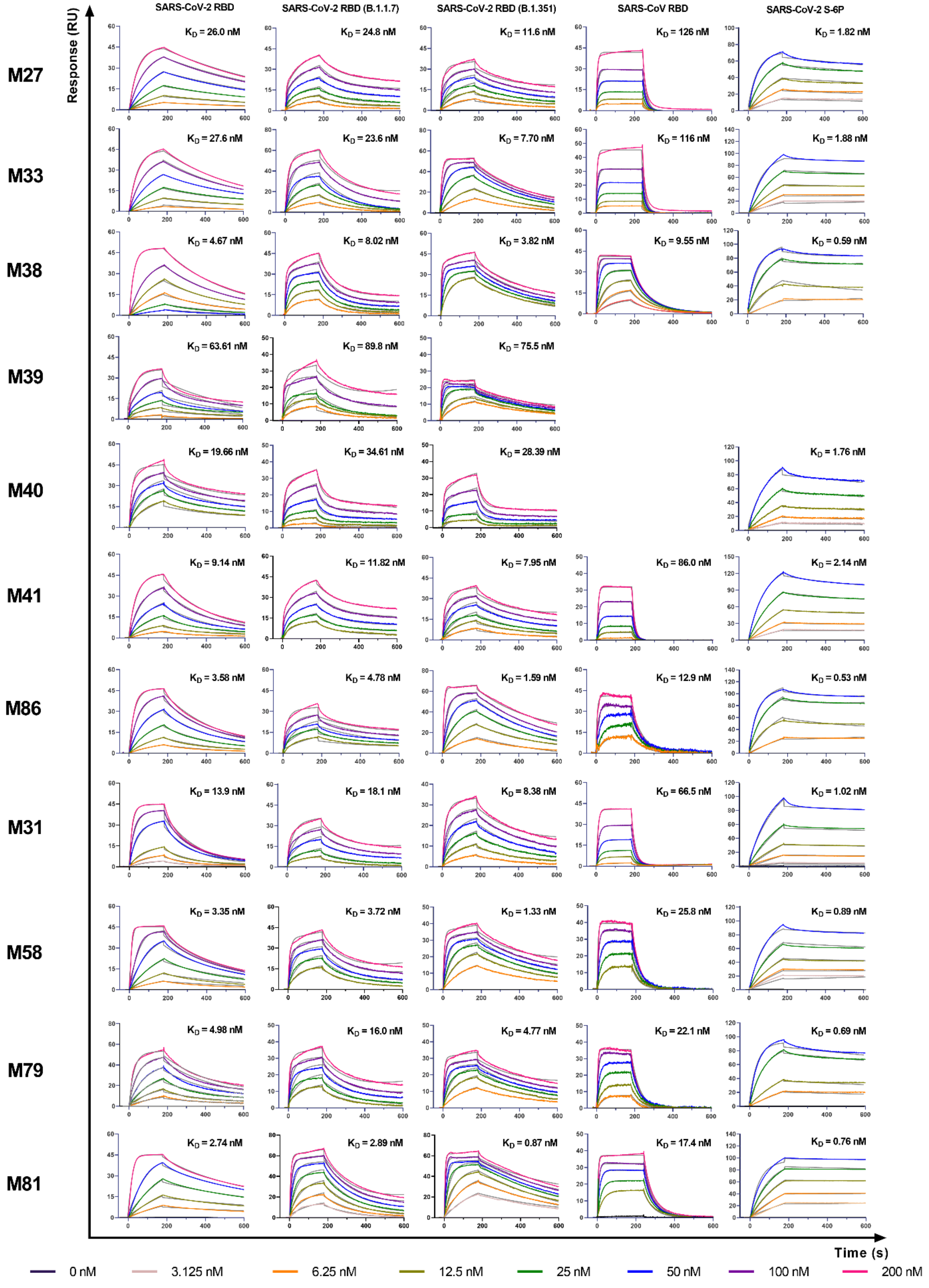

SARS-CoV-2 RBD
SARS-CoV-2 RBD (B.1.1.7)
SARS-CoV-2 RBD (B.1.351)
SARS-CoV RBD
SARS-CoV-2 S-6P
Response (RU)
Time (s)
M27
$K_D$ = 26.0 nM
$K_D$ = 24.8 nM
$K_D$ = 11.6 nM
$K_D$ = 126 nM
$K_D$ = 1.82 nM
M33
$K_D$ = 27.6 nM
$K_D$ = 23.6 nM
$K_D$ = 7.70 nM
$K_D$ = 116 nM
$K_D$ = 1.88 nM
M38
$K_D$ = 4.67 nM
$K_D$ = 8.02 nM
$K_D$ = 3.82 nM
$K_D$ = 9.55 nM
$K_D$ = 0.59 nM
M39
$K_D$ = 63.61 nM
$K_D$ = 89.8 nM
$K_D$ = 75.5 nM
M40
$K_D$ = 19.66 nM
$K_D$ = 34.61 nM
$K_D$ = 28.39 nM
$K_D$ = 1.76 nM
M41
$K_D$ = 9.14 nM
$K_D$ = 11.82 nM
$K_D$ = 7.95 nM
$K_D$ = 86.0 nM
$K_D$ = 2.14 nM
M86
$K_D$ = 3.58 nM
$K_D$ = 4.78 nM
$K_D$ = 1.59 nM
$K_D$ = 12.9 nM
$K_D$ = 0.53 nM
M31
$K_D$ = 13.9 nM
$K_D$ = 18.1 nM
$K_D$ = 8.38 nM
$K_D$ = 66.5 nM
$K_D$ = 1.02 nM
M58
$K_D$ = 3.35 nM
$K_D$ = 3.72 nM
$K_D$ = 1.33 nM
$K_D$ = 25.8 nM
$K_D$ = 0.89 nM
M79
$K_D$ = 4.98 nM
$K_D$ = 16.0 nM
$K_D$ = 4.77 nM
$K_D$ = 22.1 nM
$K_D$ = 0.69 nM
M81
$K_D$ = 2.74 nM
$K_D$ = 2.89 nM
$K_D$ = 0.87 nM
$K_D$ = 17.4 nM
$K_D$ = 0.76 nM
0 nM
3.125 nM
6.25 nM
12.5 nM
25 nM
50 nM
100 nM
200 nM

**Fig. S3. SPR kinetic measurement of SARS-CoV-2/SARS-CoV antigens binding to immobilized ACE2-Fcs. Related to Fig. 2F and Table S2.** All measurement were performed on a Protein A chip with ACE2-Fc immobilized to different levels according to the flow antigens: ~80-200 RU for SARS-CoV-2 RBD, $RBD_{B.1.1.7}$ and $RBD_{B.1.351}$, ~120 RU for SARS-CoV-1 RBD and ~60 RU for SARS-CoV-2 non-tagged S-6P. The flow antigens were injected at indicated concentrations. The background-corrected sensorgrams (colored) were fitted with 1:1 Langmuir model (grey) and the kinetic constants were summarized in **Table S1.**

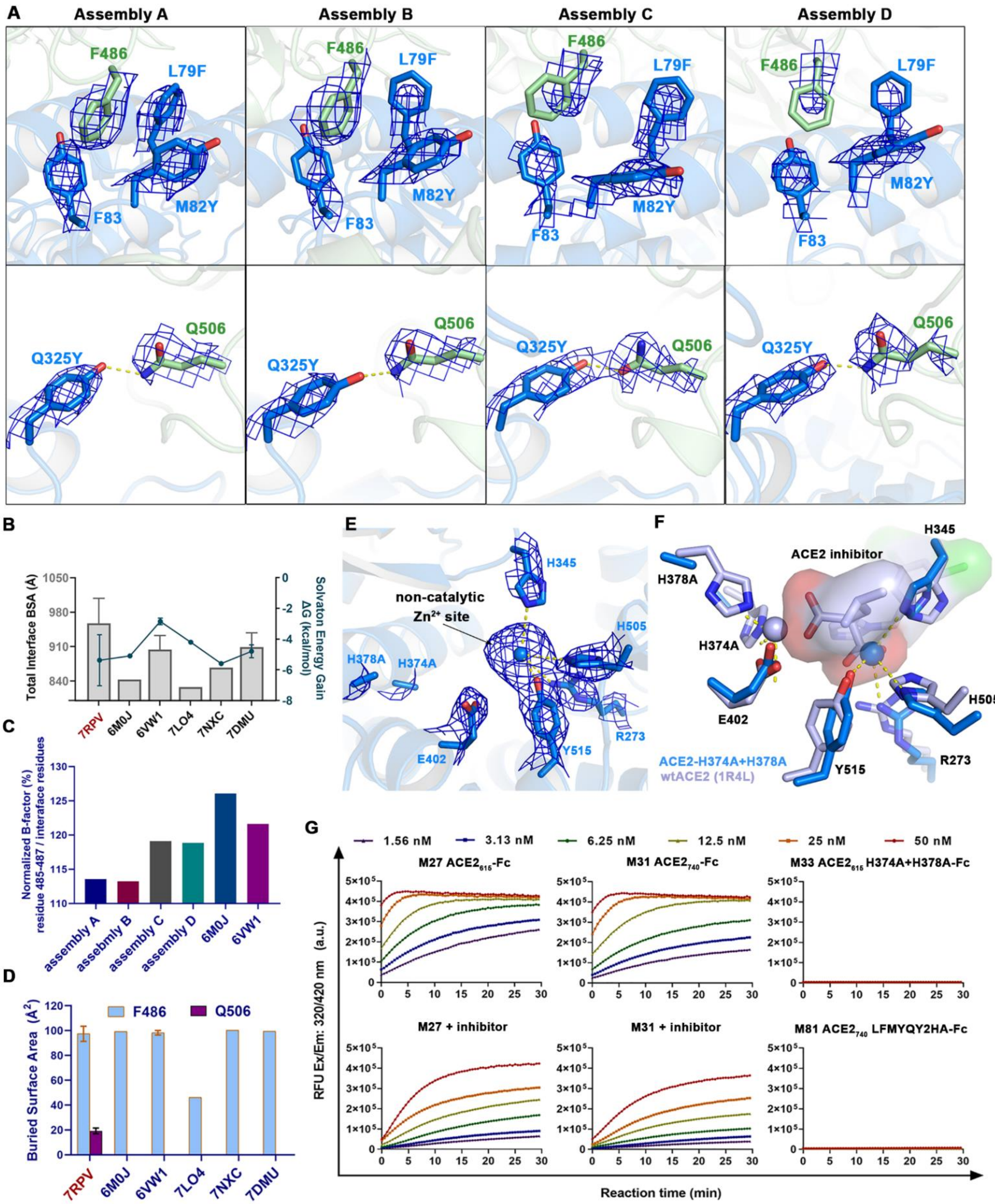

A
Assembly A
Assembly B
Assembly C
Assembly D
F486
L79F
M82Y
F83
Q325Y
Q506
B
Total Interface BSA (Å)
Solvaton Energy Gain ΔG (kcal/mol)
7RPV
6M0J
6VW1
7LO4
7NXC
7DMU
C
Normalized B-factor (%) residue 485-487 / interaface residues
assembly A
assebmly B
assembly C
assembly D
6M0J
6VW1
D
Buried Surface Area (Å²)
F486
Q506
E
non-catalytic Zn²⁺ site
H345
H505
H378A
H374A
E402
Y515
R273
F
ACE2 inhibitor
ACE2-H374A+H378A
wtACE2 (1R4L)
G
1.56 nM
3.13 nM
6.25 nM
12.5 nM
25 nM
50 nM
M27 ACE2615-Fc
M31 ACE2740-Fc
M33 ACE2615 H374A+H378A-Fc
M27 + inhibitor
M31 + inhibitor
M81 ACE2740 LFMYQY2HA-Fc
RFU Ex/Em: 320/420 nm (a.u.)
Reaction time (min)

**Fig. S4. Crystal structure of engineered $ACE2_{615}$ LFMYQY2HA in complex with SARS-CoV-2 RBD. Related to Fig. 3.** (**A**) $2F_o$-$F_c$ electron density map (contour level = 1.0 σ) of affinity-enhancing mutations L79F, M82Y and Q325Y in four assemblies within the ASU. (**B**) Total buried surface area (BSA) of -ACE2-RBD interface and solvation energy gain upon complex formation analyzed for $ACE2_{615}$ LFMYQY2HA-RBD, four wtACE2-RBD complexes (PDB: 6M0J, 6VW1, 7LO4, 7NXC) and ACE2-RBD complex where ACE2 was mutated to gain enhanced RBD affinity (PDB: 7DMU). (**C**) Normalized average B-factor of RBD ridge residues 485-487 to those of the respective ACE2-RBD interface residues among the four assemblies of $ACE2_{615}$ LFMYQY2HA-RBD structure and two wtACE2-RBD crystal structures. Interface residues were defined as those with BSA>0 $Å^2$ when calculated in PISA. (**D**) Buried surface area of two RBD residues F486 and Q506 among ACE2-RBD complexes. (**E**) $2F_o$-$F_c$ electron density map (contour level = 1.0 σ) of the active site residues in $ACE2_{615}$ LFMYQY2HA. (**F**) Active site superimposition of $ACE2_{615}$ LFMYQY2HA and wtACE2 bound to inhibitor MLN-4760. Molecular surface is displayed over the inhibitor molecule and the coordinating residues are shown as sticks. (**G**) Time course measurement of ACE2 enzyme activity. The slopes of the initial linear region were calculated and plotted as **Fig. 3G**.

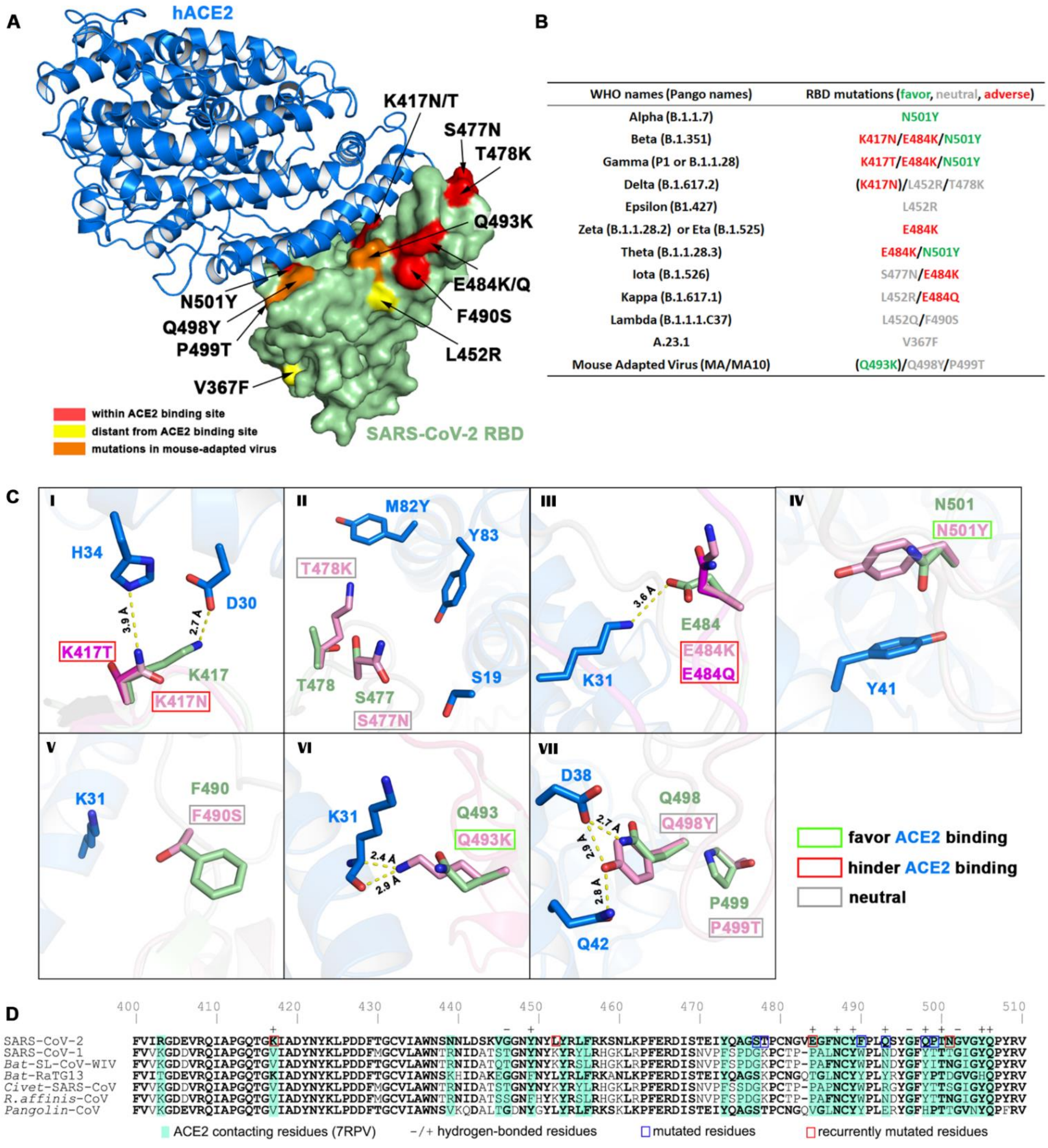

**Fig. S5. Structural basis for broad-reactivity against SARS-CoV-2 VOCs. Related to Fig. 3-4.** (**A**) Mapping of RBD mutations in SARS-CoV-2 VOCs, in the context of the $ACE2_{615}$ LFMYQY2HA-RBD structure. (**B**) Summary of RBD mutations in SARS-CoV-2 VOCs. Substitutions that were predicted to be favorable, neutral or deleterious to ACE2 binding are colored in green, grey and red respectively. (**C**)

Composite model of RBD mutations on or around the ACE2 binding site. Our structural model provide a rational explanation on the mild VOC resistance to the engineered ACE2-Fc M81. For instance, two RBD hotspot residues K417 and E484, which form salt-bridges with two ACE2 Site-I residues D30 and K31 respectively, are recurrently substituted by K417N/T and E484K/Q in several SARS-CoV-2 VOCs. These mutations abrogate the salt-bridges with ACE2 and therefore are predicted to be deleterious to RBD binding. As expected, $PsV_{B.1.526}$ harboring E484K and $PsV_{B.1.351}$ with K417N and E484K display the highest resistance to M81 ($IC_{50}$ = 2.06 and 1.14 nM respectively, **Fig. 4**) as compared to the $PsV_{D614G}$ (0.23 nM). Another frequently occurred RBD mutant N501Y, as identified in B.1.1.7, B.1.351 and P.1, is thought to enhance the receptor binding by an additional π-π interaction with $Y41_{ACE2}$ and thereby increases the viral infectivity (*6*). However, $PsV_{B.1.1.7}$ with the sole RBD mutation N501Y is less sensitive to M81 (0.52 nM) as compared to $PsV_{D614G}$ (**Fig. 4**), suggesting that prediction of ACE2-Fc cross-reactivity by only considering RBD mutation would be inadequate and mutations in N-terminal domain (NTD) and S2 subunit may affect the sensitivity to the engineered ACE2-Fcs. (**D**) Sequence alignments for receptor binding region of SARS-CoV-2 S with other five coronaviruses that utilize ACE2 as receptor. Contact residues involved in salt-bridges or H-bonds to the $ACE2_{615}$ LFMYQY2HA are marked above the sequence with (+) for the side chain and (-) for the main chain.

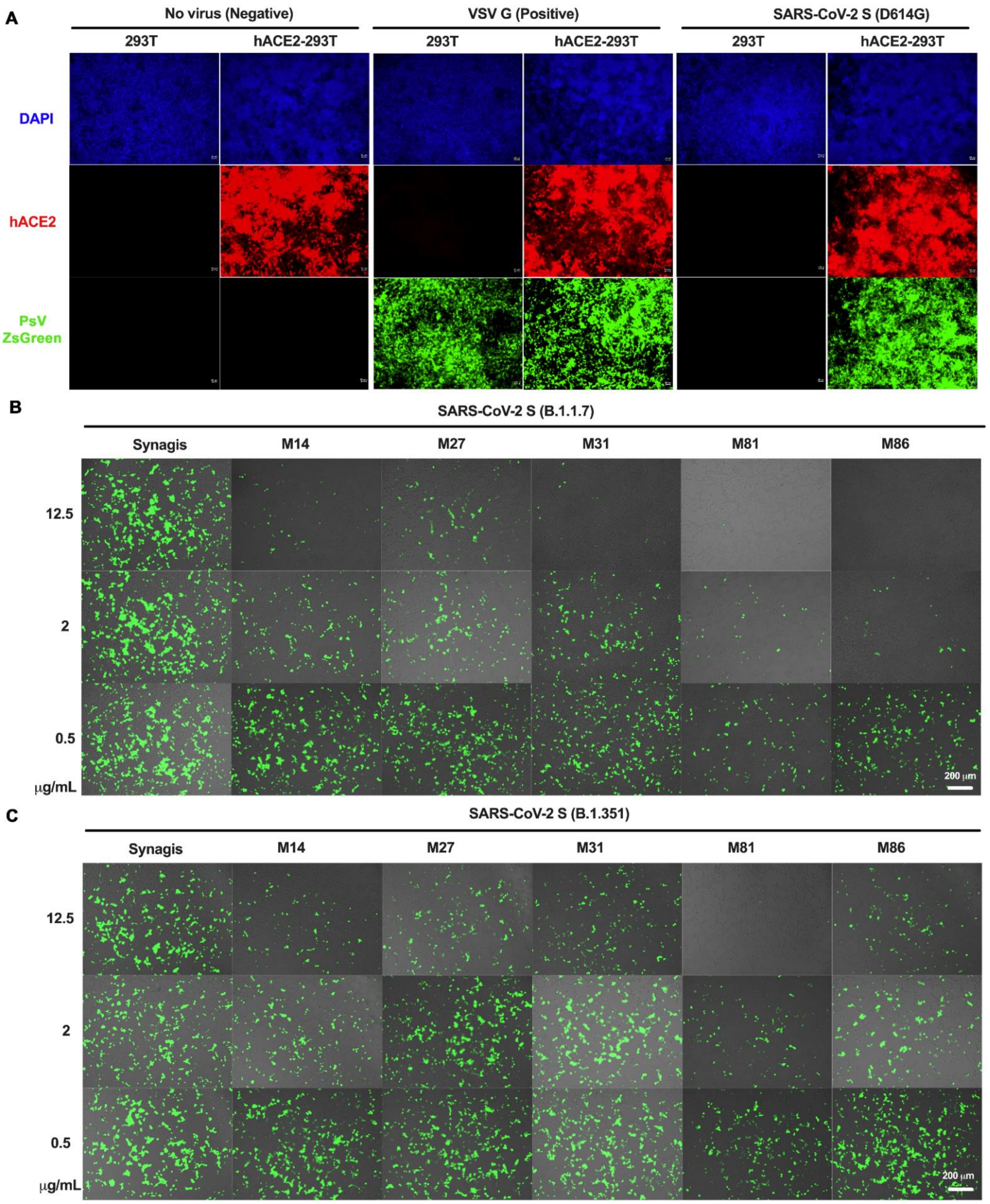

A
No virus (Negative)
VSV G (Positive)
SARS-CoV-2 S (D614G)
293T
hACE2-293T
DAPI
hACE2
PsV ZsGreen
B
SARS-CoV-2 S (B.1.1.7)
Synagis
M14
M27
M31
M81
M86
12.5
2
0.5
μg/mL
200 μm
C
SARS-CoV-2 S (B.1.351)
200 μm

**Fig. S6. Engineered ACE2-Fc variants inhibit PsV of SARS-CoV-2 VOCs from infecting hACE2-expressing 293T cells. Related to Fig. 4.** (**A**) 293T or hACE2-expressing 293T cells were treated with saline control (no virus), VSV-G PsV (positive control, ~$10^5$ RLU) or SARS-CoV-2 $PsV_{D614G}$ (~$10^6$ RLU) carrying ZsGreen reporter gene; ZsGreen signal was detected at 96 h post infection and hACE2 expression was further validated by IF staining (red) using anti-hACE2 antibody. (**B** and **C**) Representative fluorescent imaging of hACE2-expressing 293T cells that were infected with SARS-CoV-2 $PsV_{B.1.1.7}$ (**B**) or $PsV_{B.1.351}$ (**C**) in the presence of indicated concentrations of hACE2-Fc. hACE2-Fc were pre-incubated with PsV for 1 h and the protein-virus mixtures were added to hACE2-expressing 293T cells. Images taken 48h post infection were showed as merged brightfield (cell shape) and greenfield (ZsGreen signal). Scale bar: 200 μm. *n* = 3 replicates/group.

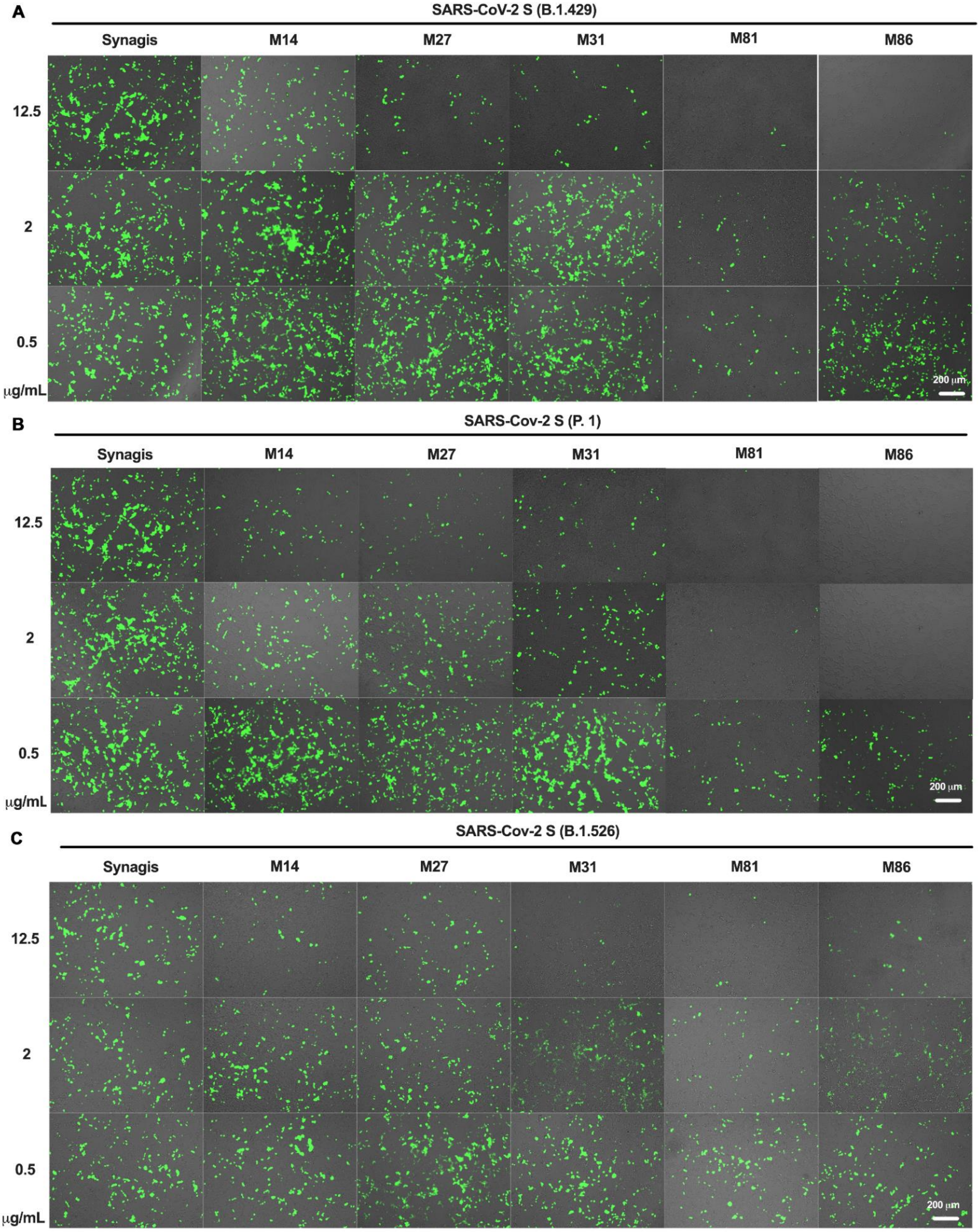

A
SARS-CoV-2 S (B.1.429)
Synagis
M14
M27
M31
M81
M86
12.5
2
0.5
μg/mL
200 μm
B
SARS-Cov-2 S (P. 1)
Synagis
M14
M27
M31
M81
M86
12.5
2
0.5
μg/mL
200 μm
C
SARS-Cov-2 S (B.1.526)
Synagis
M14
M27
M31
M81
M86
12.5
2
0.5
μg/mL
200 μm

**Fig. S7. Engineered ACE2-Fc variants inhibit PsV of SARS-CoV-2 VOCs from infecting hACE2-expressing 293T cells. Related to Fig. 4. (A-C)** Representative fluorescent imaging of hACE2-expressing 293T cells that were infected with SARS-CoV-2 $PsV_{B.1429}$ (**A**), $PsV_{P.1}$ (**B**) or $PsV_{B.1.526}$ (**C**) in the presence of indicated concentration of ACE2-Fc variants. Experimental procedures and image acquisition were identical as described in **Fig. S6**.

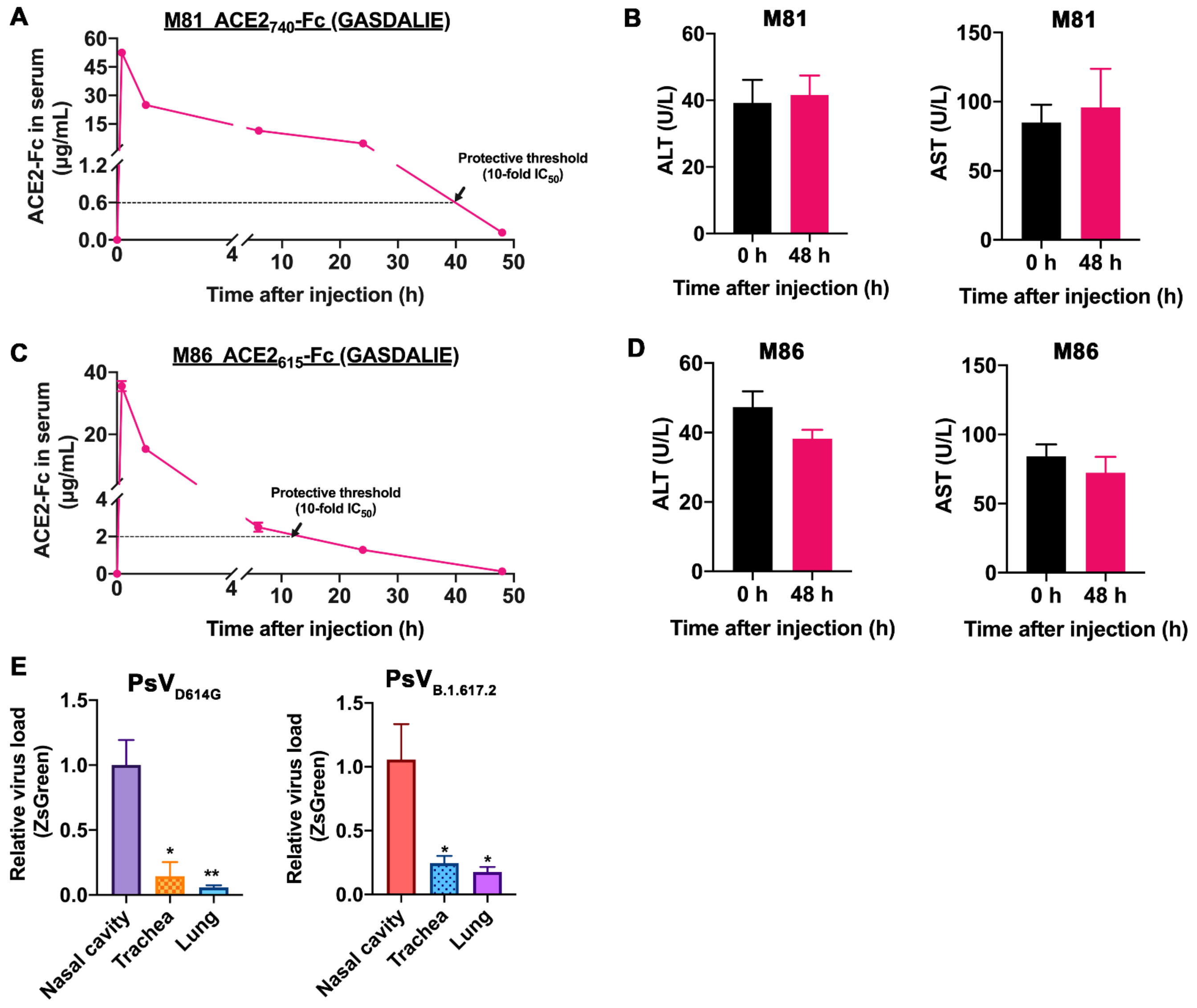

**Fig. S8. Pharmacokinetic (PK) study and hepatotoxicity test of two engineered ACE2-Fc. Related to Fig. 5.** (**A** and **C**) C57BL/6J mice were administered by 100 μg (i.v., 5 mg/kg) of two engineered ACE2-Fc variants M81 and M86. Serum samples were collected at 0 min, 10min, 1 h, 6 h, 24 h and 48 h post injection. The ACE2-Fc serum concentrations were determined by indirect ELISA (see Materials and Methods). (**B** and **D**) Serum concentrations of alanine transaminase (ALT) and aspartate transaminase (AST) before and 48 h after M81 or M86 injection. $n$ = 3 replicates/group. (**E**) The relative viral loads of nasal cavity, trachea and lung (13 dpi) in PsV-challenged K18-hACE2 mice in the Synagis-treated group (n=3-4). The relative mRNA levels of ZsGreen were calculated as fold change compared to those of nasal cavity. *$P$<0.05, versus nasal cavity.

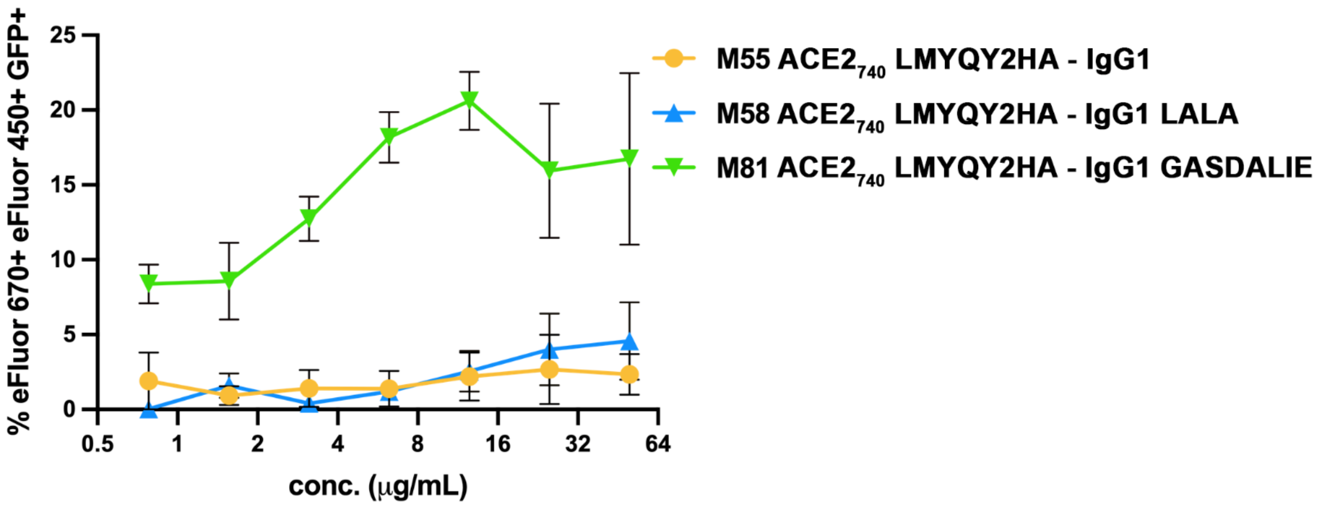

**Fig. S9. ACE2-Fc mediated phagocytosis against S-expressing CEM.NKr cells. Related to Fig. 6.** Percentage of ADCP in the presence of titrated amounts of ACE2-Fcs using CEM.NKr-Spike cells as targets and THP-1 cells as phagocytic effectors cells. Data are the average from 3 experiments; mean values ± SEM are depicted.

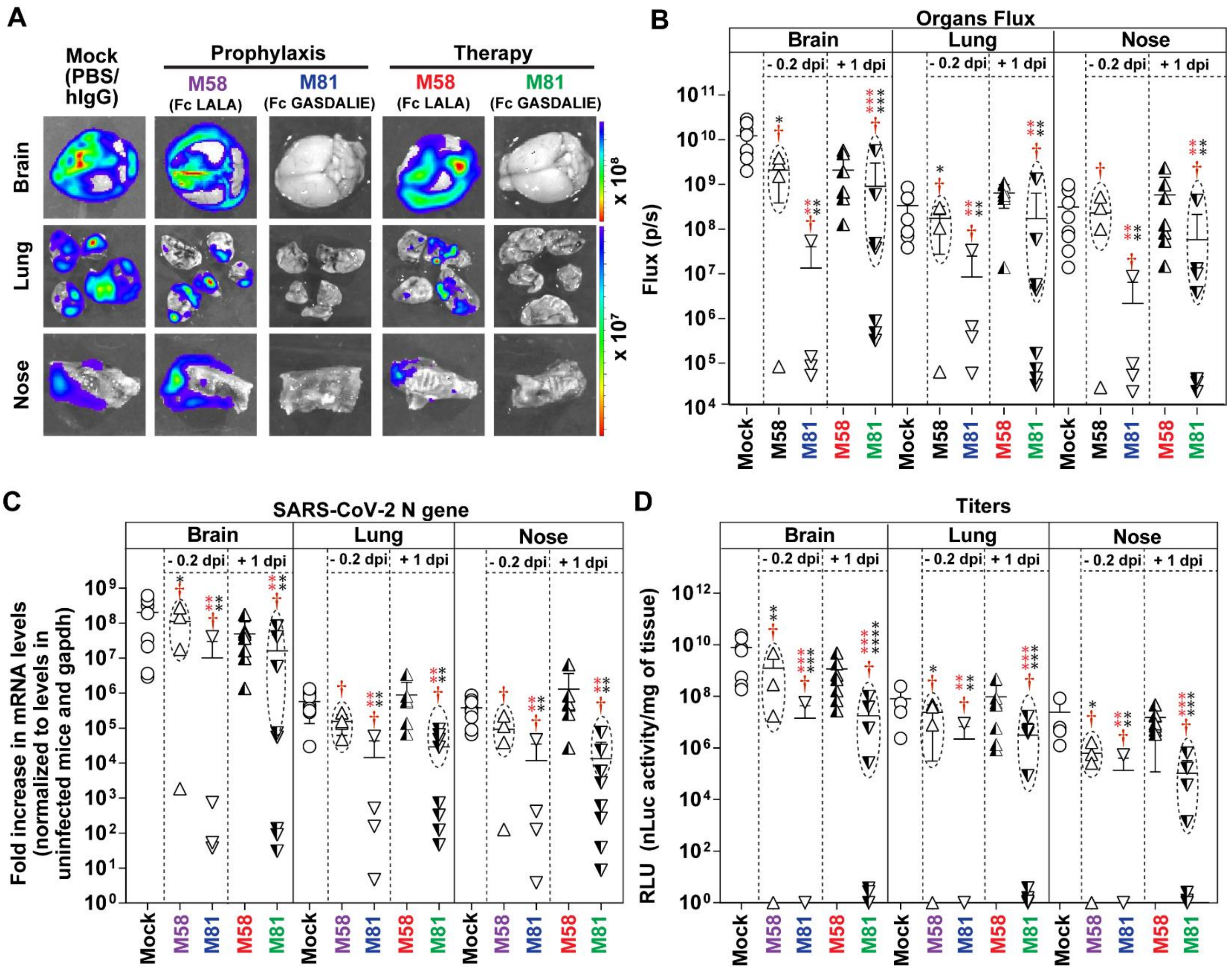

**Fig. S10. M81 significantly reduces virus loads in target organs of SARS-CoV-2-nLuc infected K18-hACE2 mice. Related to Fig. 7.** (**A-B**) *Ex vivo* imaging of indicated organs and quantification of nLuc signal as flux (photons/sec) after necropsy for an experiment shown in Figure 7. (**C**) Fold changes in nucleocapsid mRNA expression in brain, lung and nasal cavity tissues. Data were normalized to *Gapdh* mRNA in the same sample and that in non-infected mice after necropsy. (**D**) Viral loads (nLuc activity/mg) from indicated tissues using Vero E6 cells as targets. Virus loads in indicated tissues were determined when they succumbed to infection (dashed ellipse with red dagger, only for not 100% mortality cohorts) and at 20 dpi for surviving mice. Grouped data in (B-D) were analyzed by 2-way ANOVA followed by Tukey's multiple comparison tests. Statistical significance for group comparisons to control are shown in black, M58 (prophylaxis) in purple, M81 (prophylaxis) in blue, M58 (therapy) in red and M81(therapy) in green. Non-significant comparison are not shown. ∗, $P < 0.05$; ∗∗, $P < 0.01$; ∗∗∗, $P < 0.001$; Mean values ± SD are depicted.
